# CellCage™ Technology Enables High-throughput Isolation of Defined Cell Combinations to Resolve the Determinants of CAR-T mediated Cytotoxicity

**DOI:** 10.64898/2026.08.21.746043

**Authors:** Richard G Yau, Shreya Deshmukh, Makenzie Sacca, Ria Gupta, Yanping Yang, Mehdi Mohseni, Liz Y Wu, Filiz Gorpe Yasar, Shan Sabri, Teresa Ai, Tarun K Khurana, Shawn Levy, Pier Federico Gherardini, Christopher Mason, Moonsoo M. Jin, Gary P Schroth

## Abstract

Understanding the cellular determinants of CAR-T cell-mediated cytotoxicity at the single-cell level remains a critical challenge in developing effective cellular immunotherapies. Conventional bulk co-culture assays provide only population-level measurements of cytotoxic activity and cannot resolve the functional heterogeneity of individual effector-target interactions. Here, we demonstrate the Cellanome platform, which enables the formation of CellCage™ Enclosures (CCEs) through spatially controlled photopolymerization, as a tool for high-resolution, high-throughput characterization of CAR-T cell-mediated cytotoxicity. By isolating defined effector-target combinations within individual CCEs and coupling encapsulation with longitudinal time-lapse imaging and automated image analysis, we resolve cytotoxic activity across thousands of individually tracked CCEs. Using anti-CD19 CAR-T effectors and NALM6-GFP targets, we show that individual CD8^+^ CAR-T effectors exhibit substantial functional heterogeneity, with serial killing capacity, effector-target contact dynamics, and intrinsic motility each independently correlating with cytotoxic potency. Cytotoxic efficacy scaled positively with effector number, and CD4^+^ T helper cells augmented CD8^+^-mediated killing in a dose-dependent manner, providing direct evidence for cooperative T cell behavior during tumor cell clearance. Extending the platform to a solid tumor model using AIC100 anti-ICAM-1 CAR-T cells and HeLa-GFP targets, we show that individual AIC100 effectors are insufficient to mediate effective killing, while multi-effector CCEs exhibit robust cytotoxicity scaling with effector abundance and CD4^+^:CD8^+^ composition. These data establish the Cellanome platform as a broadly applicable system for dissecting the determinants of CAR-T cell cytotoxicity at single-cell resolution, with direct translational relevance to the optimization of cellular immunotherapy.

## Introduction

Chimeric antigen receptor T-cells (CAR-T cells) are autologous or allogeneic T lymphocytes engineered to express a synthetic receptor that redirects T-cell cytotoxicity toward tumor-associated antigens in an MHC-independent manner, bypassing immune evasion mechanisms exploited by many tumors^1^. Unlike conventional therapeutics, CAR-T cells function as "living drugs" capable of *in vivo* clonal expansion and long-lived immunological memory, conferring the potential for durable disease control^1^. The clinical impact has been profound: the first pediatric B-cell acute lymphoblastic leukemia (B-ALL) patient dosed with CD19-directed CAR-T cells in 2012 remains in remission today, seven products have since received FDA approval across multiple hematological malignancies, and hundreds of trials are underway exploring novel CAR designs and targets^2–4^.

Despite these achievements, a substantial proportion of patients fail to derive lasting benefit from CAR-T therapy. This underscores a critical unresolved challenge, the pronounced heterogeneity in CAR-T activity and clinical response. Inter-patient variability in T-cell subset composition and prior therapy-induced exhaustion introduces significant batch-to-batch manufacturing variability. Clinical failures due to inadequate expansion, exhaustion, or antigen escape remain common^5^. While bulk cytokine secretion has historically served as a cornerstone potency metric, *in vivo* efficacy is far more complex, depending on the CD4^+^:CD8^+^ ratio, T-cell subset composition, metabolic state, senescence marker expression, and TCR repertoire diversity^6–8^, and reliably translating *in vitro* measurements to clinical predictions remains an open challenge^9,10^.

Single-cell technologies including flow cytometry and scRNA-seq have been extensively deployed to identify pre-infusion biomarkers predictive of *in vivo* efficacy^11–14^, but are limited to phenotypic profiling and cannot directly monitor individual cell killing. Microfluidic platforms, including droplet emulsions^15–19^, microwells^20,21^, and cell-trapping devices^22,23^, have enabled the direct monitoring of individual effector-target interactions in controlled microenvironments and revealed that cytotoxic activity is strongly heterogeneous, with a small "serial killer" subpopulation responsible for the majority of target elimination^15,16,21^. However, across all of these approaches, the inability to simultaneously control encapsulation, deliver reagents dynamically, track compartments longitudinally, and link killing outcomes to precise cellular contexts in a high-throughput manner has fundamentally limited the resolution at which cytotoxic heterogeneity can be interrogated.

Here, we address these limitations using a platform that employs light-guided hydrogel polymerization to create compartments on-demand called CellCage™ Enclosures (CCEs) around cellular targets^24,25^. Permeable walls support multi-day culture and time-lapse imaging, while flexible compartmentalization logic enables parallel testing of multiple effector-to-target ratios across tens of thousands of compartments per experiment. We demonstrate how the resulting data resolve both population-level killing statistics and individual killing event dynamics, reveal cooperative killing behavior within defined effector cell groups, and apply these capabilities to both cell line models and a clinical-grade CAR-T product, establishing a scalable framework with direct applications to potency assay development, manufacturing optimization, and patient response prediction.

## Results

### Adaptive compartmentalization of cells enables the isolation of different cell populations within CCEs

CCEs are formed using a digital micromirror device that projects violet light in defined patterns around cells of interest, inducing localized photopolymerization of the surrounding hydrogel matrix to create hollow cylinders that enclose individual cells or cell groups. Unpolymerized gel is subsequently washed away, and enclosed cells can be cultured and imaged over extended periods ranging from multiple days to weeks, enabling dynamic monitoring of cellular behavior and interactions.

CCE formation supports multiple strategies that enable precise isolation of specific cell populations or defined combinations thereof. When two distinct populations are present, distinguished by differences in fluorescence or size, CCE formation can be programmed to selectively include or exclude either population, or CCEs can be formed at predetermined fixed positions for stochastic co-encapsulation of both. For targeted methods, an algorithm identifies suitable individual cells and overlays a virtual mask indicating the locations for CCE wall placement (e.g., see solid red ring in Figures 1B and 1D). Each CCE is designed to form around a single cell of interest. If another cell meeting the same criterion lies too close to allow that cell to be isolated on its own, without also enclosing the neighboring cell, CCE formation does not proceed. The user determines the CCE size (e.g., diameter of cylinder and width of the wall) and the acceptable inner and outer distances from the CCE wall that can be occupied by other cells (Figure S1).

**Figure 1.**
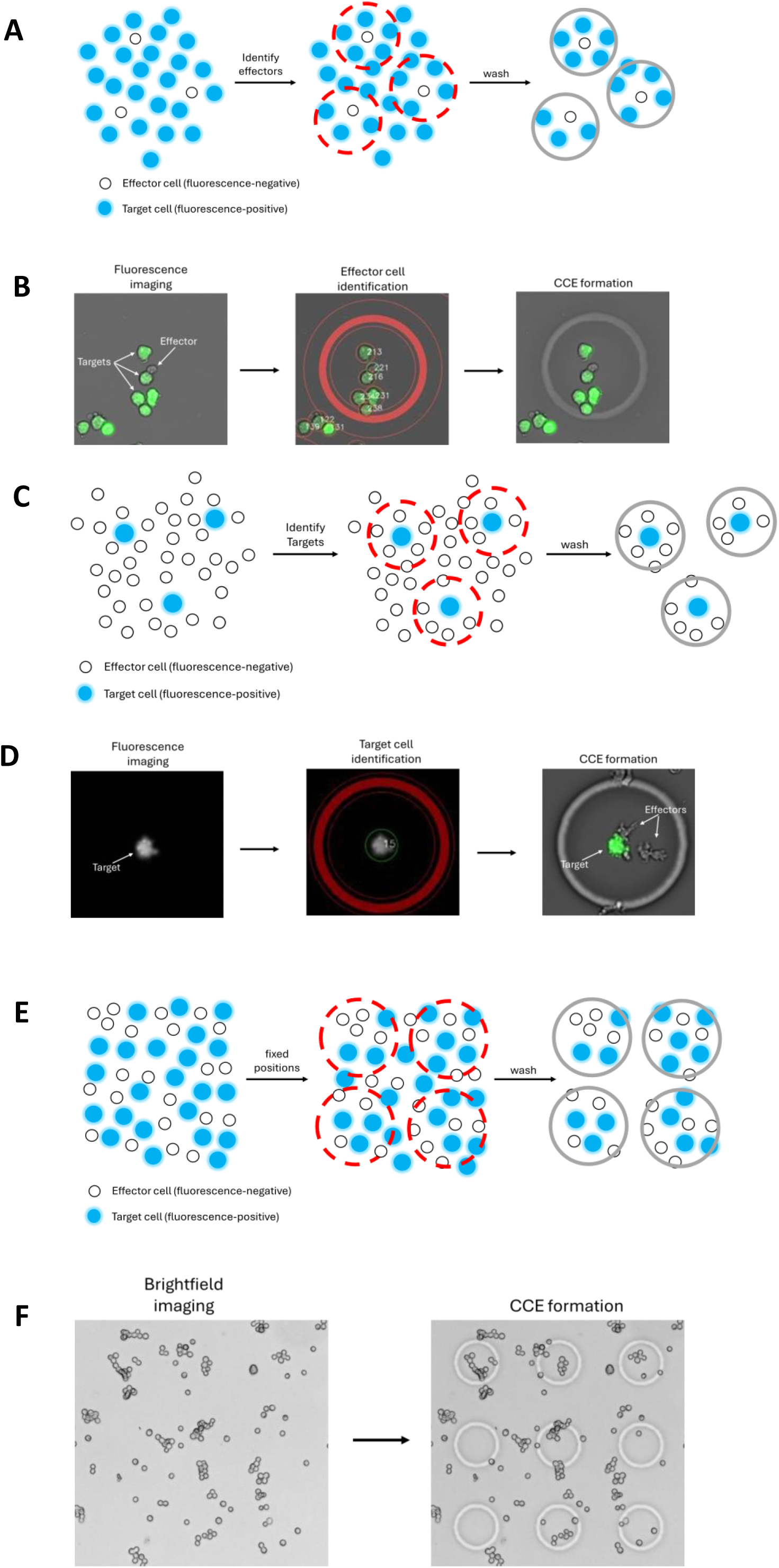
Leveraging CellCage™ technology to isolate various permutations of E:T ratios. (A) Schematic of targeted CCE formation based on identification of unfluorescent TALL-104 effector cells. CFSE-labeled K562 target cells, loaded in excess, were co-enclosed passively and in a stochastic distribution. (B) Representative CCE formed around an unfluorescent TALL-104 effector. Imaging first identified an unfluorescent cell, a mask defining the position of the CCE around the unfluorescent cell was generated, and violet light was applied to the boundary of this mask to polymerize the gel precursor forming a CCE. (C) Schematic of CCE formation based on identification of fluorescent CFSE-labeled K562 target cells. Unfluorescent TALL-104 effector cells, loaded in excess, were co-enclosed passively and in a stochastic distribution. (D) Representative CCE formed around a fluorescent target cell. Imaging first identified a fluorescent cell, a mask defining the position of the CCE around the fluorescent cell was generated, and violet light was applied to the boundary of this mask to polymerize the gel precursor forming a CCE. (E) Schematic of CCE formation based on predetermined positions, both effectors and targets were enclosed in a random manner. (F) Representative image of CCEs generated in a fixed pattern where cells were enclosed randomly.

Two targeted strategies provide bidirectional control over cell encapsulation. In the first, unfluorescent cells, TALL-104 cells were used in this example, are individually identified and selected for CCE formation, with fluorescent cells, CFSE-labeled K562 cells used here, either co- enclosed stochastically or actively excluded (Figures 1A and 1B). In the second, the targeting logic is inverted: fluorescent CFSE-labeled K562 cells are identified for CCE formation, with unfluorescent TALL-104 cells in the vicinity co-enclosed or excluded based on user-defined parameters (Figures 1C and 1D). Together, these strategies enable selective encapsulation of either population, or defined combinations of both, within a mixed sample.

CCEs can also be generated at predetermined positions across a flow cell lane without cell type identification or selection (Figures 1E-F). In this configuration, encapsulation is stochastic, with each compartment capturing cells based solely on location at the time of CCE formation, yielding a Poisson distribution of both cell populations. Because thousands of CCEs are formed without the overhead of active cell targeting, a single experiment can survey hundreds of distinct effector-to-target (E:T) ratios in parallel.

Collectively, these CCE formation strategies constitute a versatile and complementary framework for controlled cell encapsulation. The choice of method can be tailored to the experimental objective, from the precise isolation of a defined cell population to the unbiased sampling of heterogeneous cell combinations, enabling systematic interrogation of how cell-cell interactions govern biological outcomes.

### Conducting and analyzing cytotoxicity assays on the Cellanome platform

Typically, loading 1.68x10^5^ cells with 90 μm diameter CCEs generates 8,000–12,000 CCEs per lane and 64,000–96,000 per flowcell (Figure 2A-B). CCE size is user tunable, with smaller diameter CCEs producing higher throughput and larger diameter CCEs producing lower throughput. To demonstrate the platform, cytotoxicity assays were performed using CD8^+^ anti- CD19 CAR-T effectors and NALM6-GFP targets. Effectors were labeled with anti-CD8-RY586 antibodies and combined with excess NALM6-GFP targets, with GFP-negative effectors selected for CCE formation and targets passively co-enclosed (Figure S2A). Media supplemented with APC-Annexin V was applied and time-lapse imaging was performed at 6-hour intervals for 24 hours. In a representative CCE, a CAR-T effector was observed sequentially engaging multiple GFP-positive targets, which progressively acquired Annexin V fluorescence, demonstrating effector-mediated cytotoxicity at single-CCE resolution (Figures 2B and S2B).

**Figure 2.**
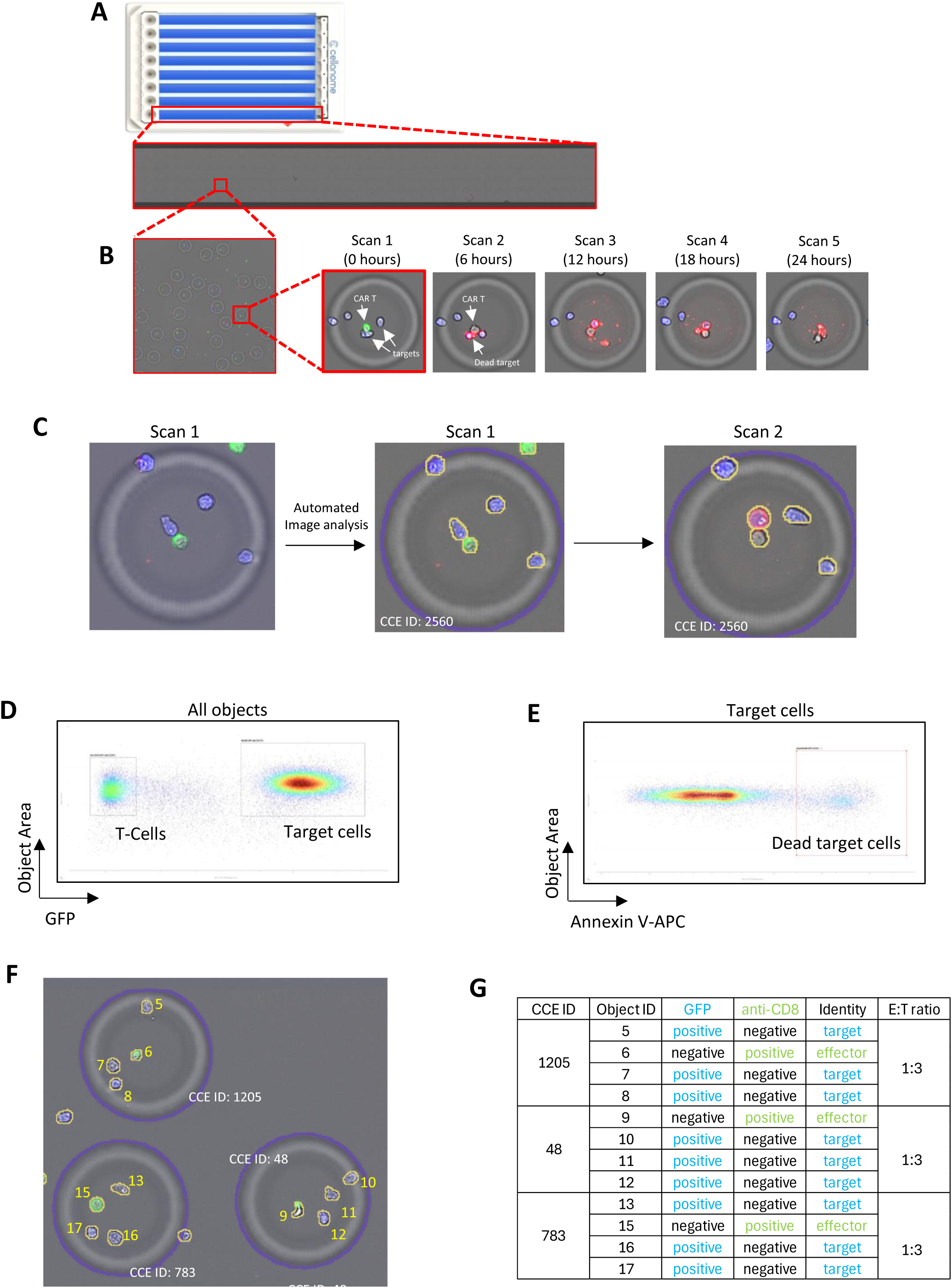
Cellanome Image analysis pipeline enables automated high-throughput quantification of cell identity in individual CCEs. (A) Each lane of an 8 lane flowcell encompasses 400 fields of view (FOV) at 10x magnification. (B) Each FOV can accommodate approximately 20-30 90 µm diameter CCEs, which translates to 8,000-12,000 CCEs per lane and 64,000-96,000 CCEs per flowcell. Cell identities were assessed based on fluorescence; NALM6-GFP cells (blue) and anti-CD8-RY586 labeled CAR-T cells (green). Cell death was monitored via change in cell number over the course of time-lapse imaging and APC-Annexin V staining (red). (C) Automated image analysis pipeline identifies all CCEs (blue outline outside the CCE) and enclosed objects (yellow outline). Every CCE and cell identified were assigned a unique identification number. (D-E) Parameters quantified for enclosed objects, including object area (size), GFP fluorescence intensity, and Annexin V-APC intensity were displayed and analyzed using Cellanome’s Cloud-based analysis platform. (D) Plotting cell size (object area) against GFP fluorescence discerns effectors from targets. (E) Gating on GFP-positive targets and interrogating for APC-Annexin V fluorescence, identifies the proportion of dead target cells. (F-G) Linking cell identities to the CCE ID in which they are enclosed. (F) Example CCEs each encapsulating a different combination of anti-CD8-RY586 labeled effectors and GFP expressing targets. Unlike object-level analyses (A– E), which treat each segmented cell as an independent observation and pool cells across CCEs, CCE-level analyses preserve the local microenvironment, analyzing each cell’s behavior in the context of the specific effectors, targets, and stoichiometry with which it shared a CCE. (G) Determination of the E:T ratio within individual CCEs. Based on fluorescence, every enclosed cell was identified as either effector or target. Cell identities were then linked back to the CCEs in which they were enclosed, and E:T ratios were determined. This method of analyses enabled the correlation of cell behaviors to specific cell combinations.

An automated imaging pipeline processes all images acquired during an experiment using an instance segmentation model to identify and uniquely label all CCEs and enclosed cells, recording positional coordinates for longitudinal CCE tracking across timepoints (Figure 2C). Cell identity is determined at each timepoint based on size (object area measured in pixels), morphology, and fluorescence. Here, effectors were resolved from targets based on GFP fluorescence (Figure 2D). Each data point is linked to the image of the cell from which it was quantified, allowing users to more accurately define gates and identify cell populations of interest (Figure S2C). GFP-negative cells were predominantly CD8-positive, based on anti-CD8-RY586 antibody labeling, confirming that these cells were the CAR-T effector population (Figure S2D). Gating on the GFP-positive target cells allowed for another gating process to be performed on this population to assess APC-Annexin V staining, thereby enabling the quantification of the proportion of dead target cells (Figure 2E). The same analysis method can be performed for each time-lapse step throughout the experiment to quantify target cell death longitudinally (Figure S2G).

### Correlating activity to cell populations within individual CCEs

Population-level analyses that aggregate data across all CCEs within a lane confirm that elevated cell death is effector-dependent, but this aggregation obscures critical biological information. Cytotoxic activity is not uniform across CCEs; rather, it depends on the specific effector-target combination present in each compartment, a dimension that aggregate readouts cannot resolve. CCE-level analysis addresses this by linking cellular identities and activities to each cell’s encapsulating compartment. Because the imaging pipeline assigns unique identifiers to both cells and CCEs, cell identity derived from population-level gating, with each cell classified as a GFP-positive/CD8-negative target or GFP-negative/CD8-positive effector, is mapped back to individual CCEs, resolving the E:T ratio for every compartment (Figures 2F and 2G). Unlike object-level analyses (Figure 2A-E), which treat each segmented cell as an independent observation in the style of conventional single-cell analysis (informative for cell-intrinsic properties but blind to interaction context), CCE-level analyses constrain cell behaviors to the individual CCEs in which they occurred, preserving the effector-target pairings, stoichiometry, and neighborhood composition that shape cytotoxic outcome. This directly links each killing event to the precise cellular context in which it occurred, transforming cytotoxicity from a population statistic into a single-event measurement and revealing compositional determinants of killing activity that aggregate approaches cannot resolve.

CCE-level analyses confer additional practical advantages. CCEs lacking targets or containing non-viable effectors are automatically excluded, reducing background signal (Figure S2E). CCE-level analyses also resolve a key confound of GFP-based cytotoxicity quantification: as dying target cells progressively lose GFP fluorescence, they become indistinguishable from GFP-negative effectors. This loss of GFP fluorescence precludes accurate target cell identification, thereby compromising population-level cytotoxic quantification, based on Annexin V, and causing underestimation of cumulative cytotoxic activity (Figures S2F-G). Because target cell number per CCE is recorded at baseline and at each time point, target cell death is quantified simply as the reduction in GFP-positive cells per CCE over time, irrespective of whether dying cells retain GFP fluorescence.

### High-resolution profiling of single cell anti-CD19 CAR-T cytotoxic activity

CD19 is the most commonly targeted antigen in CAR-T therapy, with four of seven FDA-approved products utilizing an scFv derived from the FMC63 monoclonal antibody^26–29^, which serves as a benchmark in many preclinical investigations^30–32^. To demonstrate the cytotoxicity assay using this well-established model, CD8^+^ anti-CD19 CAR-T effectors were labeled with anti- CD8-RY586 antibodies and combined with excess NALM6-GFP targets, with individual effectors identified by RY586 fluorescence for CCE formation and targets passively co-enclosed; the majority of CCEs enclosed a single effector with 0–5 targets (Figures 3A, S3A-B). A negative control lane contained NALM6-GFP targets alone. CCE formation focused on a subset of anti- CD19-RY586 labeled NALM6-GFP cells to maintain CCE formation consistency across lanes, establishing baseline cell death independent of CAR-T activity (Figure S3B).

**Figure 3.**
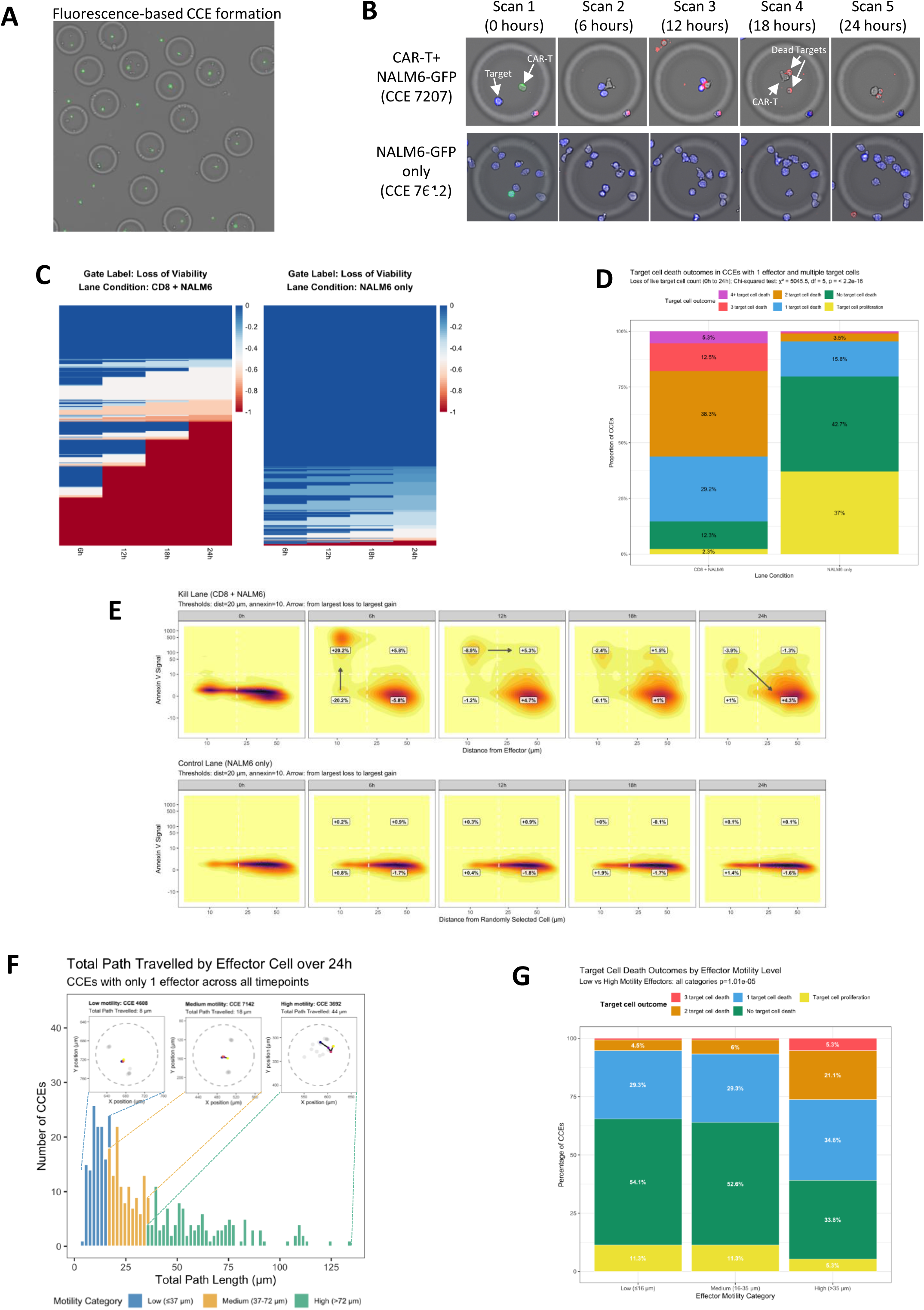
Single-cell resolution profiling of CD8^+^ CAR-T cell cytotoxic activity. (A) Fluorescence-based CCE formation identifies CD8^+^ cells, based on anti-CD8-RY586 antibody labeling, for enclosure. Unlabeled NALM6-GFP targets were co-enclosed passively. (B) Representative time-lapse brightfield and fluorescence overlay images depicting CAR-T cell-mediated cytotoxicity within individual CCEs across two conditions: a single CD8^+^ CAR-T effector co-enclosed with NALM6-GFP target cells (top row), and NALM6-GFP target cells enclosed in the absence of effectors (bottom row). Images were acquired at 6-hour intervals, with GFP fluorescence (blue) marking live target cells and APC-Annexin V fluorescence (red) indicating target cell death. (C) Longitudinal heatmaps depicting the proportion of live NALM6-GFP target cells lost since baseline within individual CCEs across two experimental conditions: CD8^+^ CAR-T effectors co-enclosed with NALM6-GFP targets (left), and NALM6-GFP targets alone (right). Each row represents a single CCE tracked across four imaging timepoints (6, 12, 18, and 24 hours), with color intensity reflecting the fraction of target cell loss from the baseline timepoint at 0 hours. (D) Stacked bar charts depicting the distribution of target cell death outcomes within individual CCEs enclosing a single CD8^+^ CAR-T effector and multiple NALM6-GFP target cells (left) compared to NALM6-GFP targets enclosed in the absence of effectors (right). The proportion of CCEs exhibiting each outcome category, ranging from target cell proliferation and no target cell death to the loss of 1, 2, 3, or 4 or more target cells, was quantified using the change in live target cell count from 0 to 24 hours as the metric. A chi-squared test was computed on the distribution of categories between the two lanes. (E) Scatter plots of individual target cell characteristics (filtered to CCEs with 1 effector cell at all timepoints to enable effector tracking), depicting the relationship between effector-target distance (x-axis, µm) and Annexin V signal of individual target cells (y-axis) across five imaging timepoints (0, 6, 12, 18, and 24 hours). Each plot is divided into four quadrants delineating proximal versus distal effector-target distances and Annexin V-positive versus Annexin V-negative target cells. The change in percentage of target cells in each quadrant is annotated, and arrows indicate the direction of the largest net population shift between quadrants at each timepoint. Spearman correlation coefficients (rho) and associated p-values are displayed for each timepoint, confirming a statistically significant negative correlation between effector-target distance and Annexin V signal across all timepoints. (F) Histogram depicting the distribution of total path lengths travelled by individual CAR-T effectors across all CCEs over a 24-hour observation period. The x-axis represents total path length (µm) and the y-axis represents the number of CCEs. Dashed vertical lines denote tertile cutoffs used to stratify effectors into low (blue), medium (yellow), and high (green) motility subpopulations. Insets show example traces of individual effectors’ paths in CCEs, with target cell positions in grey for context. The low motility CCE stays in the same position over time, the medium motility CCE moves a short overall distance, while the high motility CCE travels to a different region of the CCE, accruing a larger total path length. (G) Stacked bar charts depicting the distribution of cytotoxicity outcomes within individual CCEs stratified by effector motility category: low motility (left), medium motility (middle), and high motility (right). The proportion of CCEs exhibiting each outcome, ranging from target cell proliferation and no target cell death to the loss of 1, 2, or 3 or more target cells, is shown for each motility group. A chi-squared test was computed on the distribution of categories.

After CCE formation, cells were imaged every 6 hours for 24 hours in media supplemented with APC-Annexin V. GFP-negative effectors were observed engaging GFP-positive targets, which progressively acquired Annexin V fluorescence, while targets in the negative control remained Annexin V-negative throughout (Figure 3B). Longitudinal tracking confirmed progressive, effector-dependent killing detectable from the earliest timepoints, with a substantial proportion of CCEs reaching peak killing by 18–24 hours and heterogeneous killing kinetics across the CCE population (Figures 3C and S3C). We fit a linear mixed-effects (LME) model to the data using restricted maximum likelihood (REML), with the rate of target cell death as the response variable, time as a continuous fixed effect, and the lane as a categorical fixed effect. In the presence of CD8^+^ cells, target cells died faster than in the NALM6-GFP only control lane, with a p-value < 0.001 using the Type III ANOVA test.

CCE-level analysis of GFP-positive target cell number over time demonstrated substantial serial killing capacity: the majority of effector-containing CCEs exhibited at least one target cell death event, with 29.2%, 38.3%, 12.5%, and 5.3% displaying loss of 1, 2, 3, or ≥4 targets, respectively, compared to 15.8%, 3.5%, and 1.1% in the negative control (chi-squared p < 0.001). In the control lane, target cell death was observed in 20.4% of CCEs, 42.7% of CCEs showed no target cell death, and in 37.0% of CCEs target cell proliferation was observed reflecting their active expansion in the absence of cytotoxic pressure. By contrast, at least 1 target cell died in 85.3% of CCEs with CAR-T effectors, representing a 4-fold increase in target cell death compared to the control (Figure 3D). Together, these data demonstrate that a subset of CD8^+^ CAR-T effectors is capable of serial killing. The Cellanome platform resolved and quantified multiple target killing events at single-CCE resolution, making this platform well suited to characterize the functional heterogeneity of cell therapy products. Effector-target proximity, the distance between an unfluorescent effector and fluorescent target, in µm, was measured at each time point for each CCE. To determine the relationship between effector-target proximity and target cell death, Annexin V staining (y-axis) was plotted against proximity to an effector (x-axis). As expected, live targets in contact with an effector at the initial time point (lower-left quadrant), were more likely to undergo cell death at later time points (upper left quadrant). Target cell death peaked at 6 hours when 20.2% of effector-proximal targets acquired Annexin V fluorescence. Interestingly, after the 6 hour time point, effector-target distances progressively increased, consistent with effectors disengaging from killed targets and migrating away (Figure 3E and S3D). Conversely, 59.8% of target cells were spatially distant from effectors at the initial time point (lower right quadrant) and these targets remained Annexin V-negative throughout the experiment. Using the same analysis method for the control lane, we measured the distance of target cells to a randomly nominated single cell in the same CCE. In the absence of CAR-T effectors, target cells predominantly well separated from each other and showed low Annexin V uptake at all timepoints. The distribution of cells into quadrants based on distance and Annexin V signal was significantly different between the two conditions using a chi-squared test at each timepoint.

Given that cytotoxicity is driven in part by physical effector-target proximity, we hypothesized that highly motile effectors would more effectively engage with and kill targets. The imaging pipeline records the position of each object segmented within each CCE. Given that each CCE contained a single unfluorescent effector, it was possible to track the position of individual effectors across time points and from that, calculate the path length travelled in µm over the course of the experiment. Effectors, stratified into low, medium, and high motility groups based on total path length over the 24 hour experiment showed markedly distinct cytotoxic profiles (Figure 3F and S3E). High motility effectors were associated with both greater killing activity overall, and with serial killing (the sequential engagement of multiple targets within a single CCE). CCEs in the high-motility group were more likely to show loss of 2 targets (21.1% vs. 4.5% in the low motility group) or ≥3 targets (5.3% vs. 0.9%), and were less likely to show no target cell death at all (33.8% vs. 54.1%; chi-squared p = 0.000011; Figure 3G). Together, these results reveal substantial functional heterogeneity within CAR-T cell populations, with serial killing capacity, effector-target contact dynamics, and intrinsic motility each correlating with cytotoxic potency, a level of resolution fundamentally inaccessible to conventional population-level assays.

### Cooperation between multiple CD8^+^ Cytotoxic T lymphocytes leads to highly efficient cytotoxicity

While individual anti-CD19 CAR-T effectors exhibited cytotoxic activity, studies suggest that killing is most effective through the cumulative cytolytic activities of multiple Cytotoxic T lymphocytes (CTLs)^33,34^, coordinated in part through CTL swarming behavior^35^. To test this, fixed pattern CCE formation was used to co-enclose multiple CD8^+^ CAR-T effectors with NALM6-GFP targets across a broad range of E:T ratios, with CCEs containing 0–3 effectors and 0–5 targets most abundantly represented and individual E:T combination counts reaching up to 843 CCEs (Figures 4A-C). The co-culture lane included over 1,700 effector-free CCEs serving as an internal negative control, while the NALM6-GFP-only lane predominantly enclosed 3–4 targets per CCE (Figure S4B). This compositional diversity enables systematic, high-throughput analysis of how E:T ratio influences cytotoxic outcomes across thousands of individually tracked enclosures.

**Figure 4.**
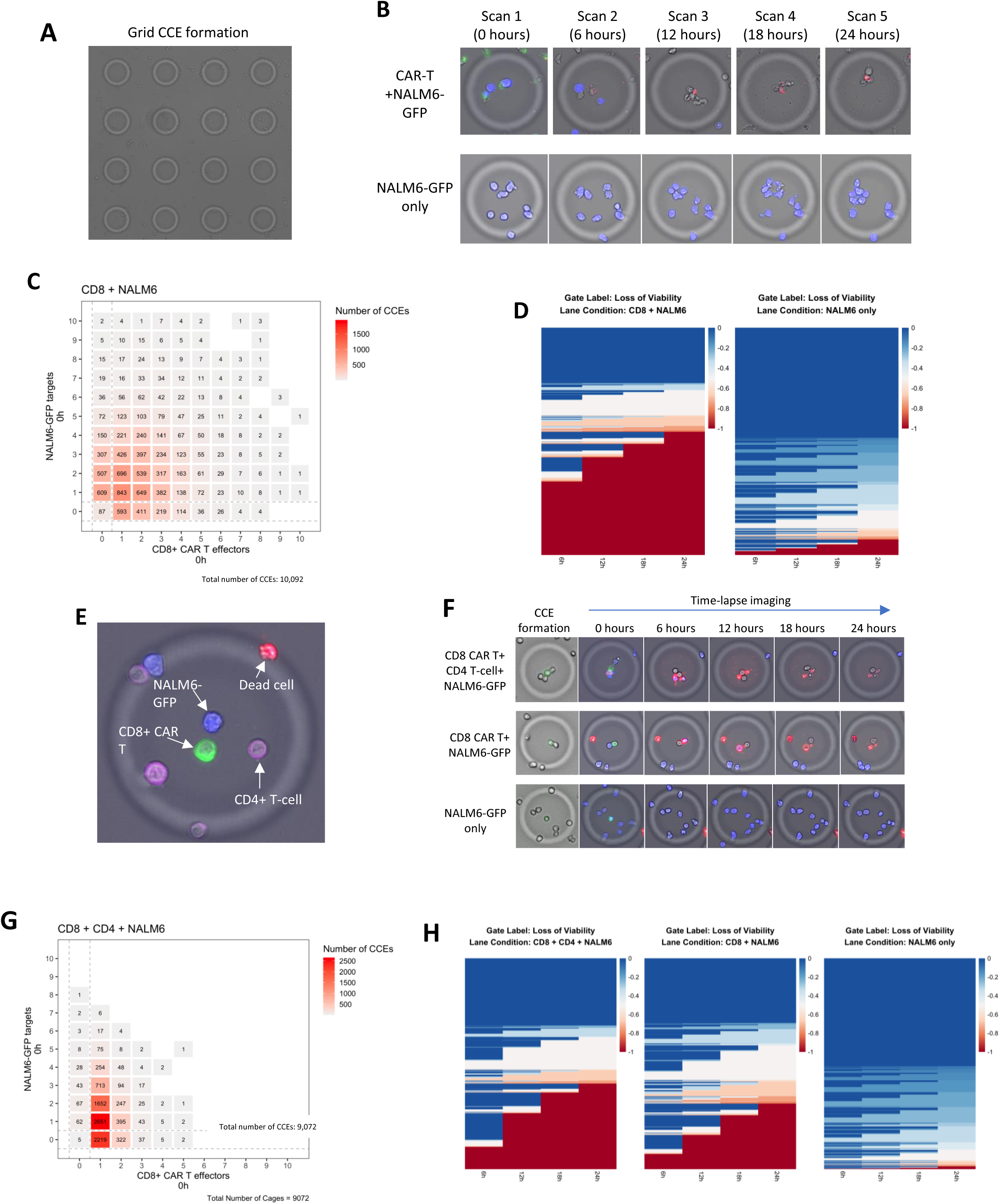
Cooperative and dose-dependent CAR-T cell-mediated cytotoxicity resolved at single-CCE resolution using fixed pattern encapsulation. (A) Fixed pattern CCE formation arranged in a grid incorporates a stochastic distribution of effectors and targets. Cell identities were not assessed during CCE formation as encapsulation was passive for both effectors and targets. (B) Representative time-lapse images depicting cytotoxic activity within individual CCEs across two conditions: multiple CD8^+^ CAR-T effectors co-enclosed with NALM6-GFP target cells (top row), and NALM6-GFP target cells enclosed in the absence of effectors (bottom row). Images were acquired at 6-hour intervals over 24 hours. GFP fluorescence (blue) marks live NALM6-GFP target cells, green fluorescence identifies CAR-T effectors, and APC-Annexin V fluorescence (red) indicates target cell death. (C) Heatmap depicting the distribution of CCE compositions at baseline (0 hours), with the number of CD8^+^ CAR-T effectors (x-axis) and NALM6-GFP target cells (y-axis) per CCE shown across a total of 10,092 CCEs. Each tile represents a unique effector-target combination, with color intensity and annotated numerical values reflecting the number of CCEs containing that specific composition. (D) Longitudinal heatmaps displaying the proportion of live target cells lost since baseline across individual CCEs, with each row representing a single CCE and each column corresponding to a time point (6, 12, 18, and 24 hours). The color scale reflects the fraction of live target cell loss from the baseline timepoint at 0 hours within each CCE, where blue indicates no target cell death and red indicates complete target cell death. Left; NALM6-GFP targets co-enclosed with CD8^+^ CAR-T effectors. Right; NALM6-GFP targets alone. (E) Representative CCE where individual anti-CD8-RY586 labeled CD8^+^ CAR-T effectors were identified for fluorescence-based CCE formation. Anti-CD4-Alexa Fluor 405 labeled CD4^+^ T lymphocytes and GFP-positive target cells were passively co-enclosed. (F) Representative time- lapse images depicting CAR-T mediated cytotoxic activity within individual CCEs shown over the indicated imaging timepoints. Top row; CCE encapsulating CD8^+^ CAR-T, CD4^+^ untransduced T lymphocytes, and NALM6-GFP target. Middle row; CCE encapsulating CD8^+^ CAR-T and NALM6-GFP in the absence of CD4^+^ untransduced T lymphocytes. Bottom row; CCE encapsulating NALM6-GFP alone in the absence of CD8^+^ CAR-T effectors and CD4^+^ untransduced T lymphocytes. (G) Heatmap depicting the distribution of CCE compositions at baseline (0 hours), with the number of CD8^+^ CAR-T effectors (x-axis) and NALM6-GFP target cells (y-axis) per CCE shown across a total of 9,072 CCEs. Each tile represents a unique effector- target combination, with color intensity and annotated numerical values reflecting the number of CCEs containing that specific composition. (H) Longitudinal heatmaps depicting the proportion of NALM6-GFP target cell viability loss over time within individual CCEs across three experimental conditions: CD8^+^ CAR-T effectors co-enclosed with CD4^+^ untransduced T cells and NALM6-GFP targets (left), CD8^+^ CAR-T effectors co-enclosed with NALM6-GFP targets (middle), and NALM6-GFP targets alone (right). Each row represents a single CCE tracked across four imaging timepoints (6, 12, 18, and 24 hours), with color intensity reflecting the fraction of live target cell loss since baseline at 0 hours (blue, minimal loss; red, maximal loss).

After CCE formation, media supplemented with APC-Annexin V was applied and images were acquired at 6-hour intervals for 24 hours, with anti-CD8-RY586 fluorescence imaged at the first timepoint only to record effector numbers per CCE, and GFP and APC-Annexin V fluorescence imaged throughout (Figures 4B and S4A). In multi-effector CCEs, target cells progressively acquired Annexin V fluorescence with concurrent loss of GFP signal, with 74.5% of targets categorized as dead by the final timepoint. In contrast, target cells in the negative control retained GFP fluorescence throughout and expanded over time, consistent with active proliferation in the absence of cytotoxic pressure (Figures 4B and 4D). Longitudinal tracking confirmed that a large proportion of effector-containing CCEs reached peak killing within the first 6 hours. Fitting a linear mixed-effects model again showed that target cells lost viability faster in the presence of effectors than in the control lane (p-value <0.001; Figure 4D). Killing magnitude, defined as the percent of target cell death, scaled positively with effector number per CCE and was highest in the 0–6 hour window (Figures S4C–D), demonstrating that multi-CTL cytotoxicity is both rapid and dose-dependent, with the strongest effect occurring immediately after effector-target contact.

### Testing cooperativity between co-enclosed CD4^+^ T-helper and CD8^+^ Cytotoxic T lymphocytes

While the role of CD4^+^ T helper cells in priming CD8^+^ CTL responses is well established, their contribution during the effector phase remains incompletely understood^36,37^. To interrogate CD4^+^-CD8^+^ cooperativity directly, CD8^+^ anti-CD19 CAR-T effectors were co-enclosed with untransduced donor-matched CD4^+^ T cells and NALM6-GFP targets. As CD4^+^ T cells do not express the anti-CD19 CAR and cannot directly recognize targets, any enhancement in killing reflects indirect cooperative effects. CD8^+^ and CD4^+^ cells were labeled with anti-CD8-RY586 and anti-CD4-Alexa Fluor 405 antibodies, with CD8^+^ effectors selected for CCE formation and CD4^+^ and NALM6-GFP cells passively co-enclosed (Figure 4E). Media supplemented with APC-Annexin V was applied and time-lapse images were acquired at 6 hour intervals for 24 hours. Anti- CD8-RY586 and anti-CD4-Alexa Fluor 405 fluorescence were imaged in the first timepoint while GFP and APC-Annexin V fluorescence were imaged in every timepoint (Figure S4E). The majority of CCEs contained 1 CD8^+^ effector, 0-6 CD4^+^ T cells, and 0-5 NALM6-GFP targets (Figures 4G and S4F). Two control conditions were included in parallel, CD8^+^ effectors co-enclosed with targets in the absence of CD4^+^ T cells, and targets enclosed without any T cells as a negative control (Figure S4F).

In the presence of CD8^+^ CAR-T effectors or CD8^+^ CAR-T effectors with CD4^+^ T-cells, target cells progressively acquired Annexin V fluorescence, while targets enclosed without effectors maintained high viability throughout the experiment (Figure 4F). Longitudinal CCE-level tracking revealed that CD8^+^ CAR-T mediated cytotoxicity, which reached maximum levels by 18-24 hours, was further enhanced in the presence of CD4^+^ T helper cells, with a greater proportion of CCEs displaying earlier onset and more complete target cell elimination compared to the CD8^+^-only condition (Figure 4H). Target cell death scaled positively with CD4^+^ cell number per CCE (Figure S4G), demonstrating a dose-dependent cooperative effect. Collectively, these data demonstrate that CD4^+^ lymphocytes augment CD8^+^ CAR-T cytotoxicity at the single- CCE level, a cooperative behavior that would be obscured in bulk co-culture assays.

### Establishing a stain-free cytotoxicity assay using a solid tumor cell model

To further demonstrate platform versatility, cytotoxicity assays were extended to a solid tumor model using AIC100 CAR-T cells which target ICAM-1 expressing malignant thyroid cancers. HeLa cells, which express endogenous ICAM-1, were engineered to express GFP to serve as targets for this killing model^38,39^. GFP-negative AIC100 effectors were labeled with anti-CD4- RY586 and anti-CD8-Alexa Fluor 405 antibodies and combined with HeLa-GFP targets. Individual effectors were then selected for CCE formation and HeLa-GFP targets were passively co-enclosed (Figures 5A and S5A). The majority of CCEs contained 1 AIC100 effector and 0–4 HeLa-GFP targets, with 2,572 and 839 CCEs containing a single CD4^+^ or CD8^+^ effector, respectively (Figures 5B, S5B). As a negative control, HeLa-GFP targets alone were enclosed without CAR-T effectors. Here, CCEs predominantly contained 0-6 HeLa-GFP targets (Figure S5C). After CCE formation, cells were incubated in media and imaged every 4 hours over 24 hours. As HeLa-GFP targets are intrinsically fluorescent and morphologically distinct from CAR-T effectors, we hypothesized that target cell death could be quantified by loss of GFP fluorescence and changes in cell morphology, a stain-free readout supported by the automated imaging pipeline’s quantification of fluorescence and morphology (as measured by eccentricity and size) during cell segmentation. HeLa-GFP targets were readily discernible from GFP-negative CAR-T effectors (Figure S5D), and were further resolved into GFP-high/large (live) and GFP-low/small (dead) subpopulations based on fluorescence intensity and cell size. At 0 hours, GFP-high targets, and GFP-low targets accounted for 89.86% and 9.83% of GFP-positive cells, respectively; by 24 hours, the GFP-low population expanded modestly to 14.32% with a corresponding reduction in GFP-high cells to 85.16%, indicating minimal cytotoxic activity by single effectors (Figures 5C– D). Longitudinal analysis of 6,009 CCEs revealed that target cell death in single-effector CCEs was indistinguishable from the negative control, with effector-target interactions rarely resulting in cytotoxicity (Figure 5E and 5F). Furthermore, no correlation was observed between target cell death and effector proximity (Figure S5E), indicating that individual AIC100 effectors are insufficient for effective HeLa-GFP killing.

**Figure 5.**
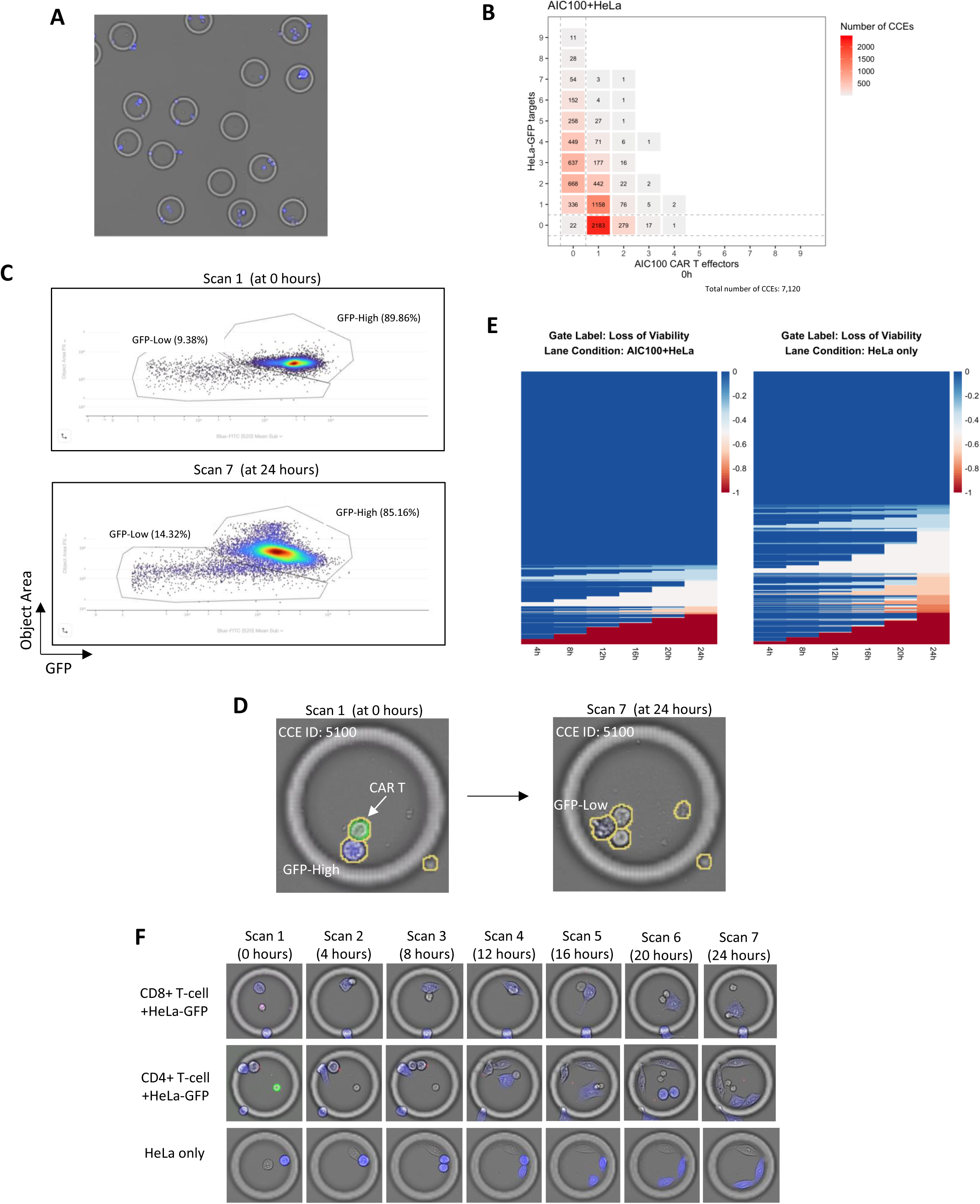
Stain-free profiling of cytotoxic activity of individual anti-ICAM AIC100 CAR-T effectors against solid tumor targets. (A) Subtractive CCE formation identifies GFP-negative AIC100 CAR-T effectors for encapsulation. HeLa-GFP targets were co-enclosed passively. (B) Heatmap depicting the distribution of CCE compositions at baseline (0 hours) for the AIC100+HeLa condition, with the number of live HeLa-GFP target cells (y-axis) and live AIC100 CAR-T effectors (x-axis) per CCE shown across a total of 7,120 CCEs. Each tile represents a unique effector-target combination, with color intensity and annotated numerical values reflecting the number of CCEs containing that specific composition. (C) Dot plots depicting the distribution of enclosed cells within CCEs based on GFP fluorescence intensity (x-axis) and cell size (y-axis) at two timepoints: 0 hours (top) and 24 hours (bottom). Two target cell populations are resolved by gating: HeLa-GFP target cells (GFP-high/large), and dead HeLa-GFP target cells (GFP-low/small). Each data point represents an individual cell. (D) Representative images of a single CCE (ID: 5100) at two timepoints, illustrating the transition of a HeLa-GFP target cell from the GFP-high/large to the GFP-low/small gate over the course of the experiment. Anti-CD4 (green) antibody fluorescence identified CD4^+^ CAR-T effectors at the initial timepoint only. At the initial timepoint (top), a GFP-high/large HeLa-GFP target cell (blue) is clearly identifiable within the CCE, exhibiting strong GFP fluorescence and a large cell area consistent with a viable target cell. By the final timepoint (bottom), the previously GFP-high target cell has undergone a marked reduction in GFP fluorescence intensity and cell size, transitioning to the GFP-low/small gate, indicating cell death has occurred. (E) Longitudinal heatmaps depicting the proportion of live HeLa-GFP target cells lost over time within individual CCEs across two experimental conditions: AIC100 CAR-T effectors co-enclosed with HeLa-GFP targets (left), and HeLa-GFP targets enclosed in the absence of effectors (right). Each row represents the loss of target cell viability from the initial timepoint at 0 hours for single CCEs tracked across multiple imaging timepoints (4 through 24 hours), with color intensity reflecting the magnitude of target cell viability loss, ranging from no loss (blue) to complete target cell elimination (red). (F) Representative time-lapse images depicting the behavior of individual AIC100 CAR-T effectors co-enclosed with HeLa-GFP target cells, alongside a negative control with only HeLa-GFP targets, over seven successive imaging timepoints. Anti-CD8 (violet) and anti-CD4 (green) antibody fluorescence identified CD8^+^ and CD4^+^ CAR-T effectors, respectively, at the initial timepoint only. GFP fluorescence (blue) was imaged in all timepoints to identify HeLa-GFP targets. Despite physical interactions between the effector and targets being evident at early timepoints, target cells largely retain their GFP fluorescence and morphology across successive imaging timepoints, indicating that the individual effector failed to mediate effective cytotoxicity.

Consistent with prior studies demonstrating that effective tumor cell killing requires the concerted activity of multiple CAR-T cells^33,34^, individual AIC100 effectors failed to kill HeLa-GFP targets. Fixed pattern CCE formation was therefore used to co-enclose multiple AIC100 effectors with HeLa-GFP targets across a broad range of compositions (Figure S6A). Effector and target numbers ranged from 0 to 16 per CCE and the most abundantly represented combinations occurring at 3-8 effectors and 0-5 targets (Figures 6A-B). A negative control lane contained HeLa-GFP targets alone, with most CCEs encapsulating 1-12 targets (Figure S6B). Targets were resolved from effectors using GFP fluorescence and further stratified into GFP-high/large and GFP- low/small subpopulations (Figures 6C-D, and S6C). Co-encapsulation of multiple AIC100 effectors significantly increased the proportion of targets transitioning from GFP-high/large to GFP-low/small over 24 hours. At 0 hours, GFP-high targets, and GFP-low targets accounted for 76.70% and 20.97% of GFP-positive cells, respectively; by 24 hours, the GFP-low population expanded to 68.46% with a corresponding reduction in GFP-high cells to 28.52%, indicating significant cytotoxic activity when multiple effectors were co-enclosed with targets (Figures 6C– D). A high proportion of effector-target interactions resulted in target cell death (Figures 6C and 6E). Longitudinal CCE-level tracking revealed progressive and widespread killing detectable from 6 hours, with heterogeneous killing kinetics across the CCE population consistent with stochastic effector-target encounter dynamics. CCEs in the HeLa-GFP-only control maintained high target viability throughout, confirming effector-dependent cytotoxicity. Fitting a linear mixed-effects model to these temporal trends again showed a significantly higher rate of target viability loss in the presence of AIC100 CAR-T effectors (p-value <0.001) (Figure 6F). Further analysis of individual CCE compositions revealed that CCEs with high E:T ratios exhibited significantly greater target cell death compared to those with low E:T ratios, consistent with the dose-dependent nature of effector-mediated killing (Figure S6D). Performing a Wilcoxon test on paired groups of CCEs with the same number of HeLa-GFP targets between lanes, the presence of AIC100 CAR-T effectors and targets was associated with a significantly greater loss of viability than CCEs with only targets, and this effect was stronger at higher E:T ratios (Figure S6D).

**Figure 6.**
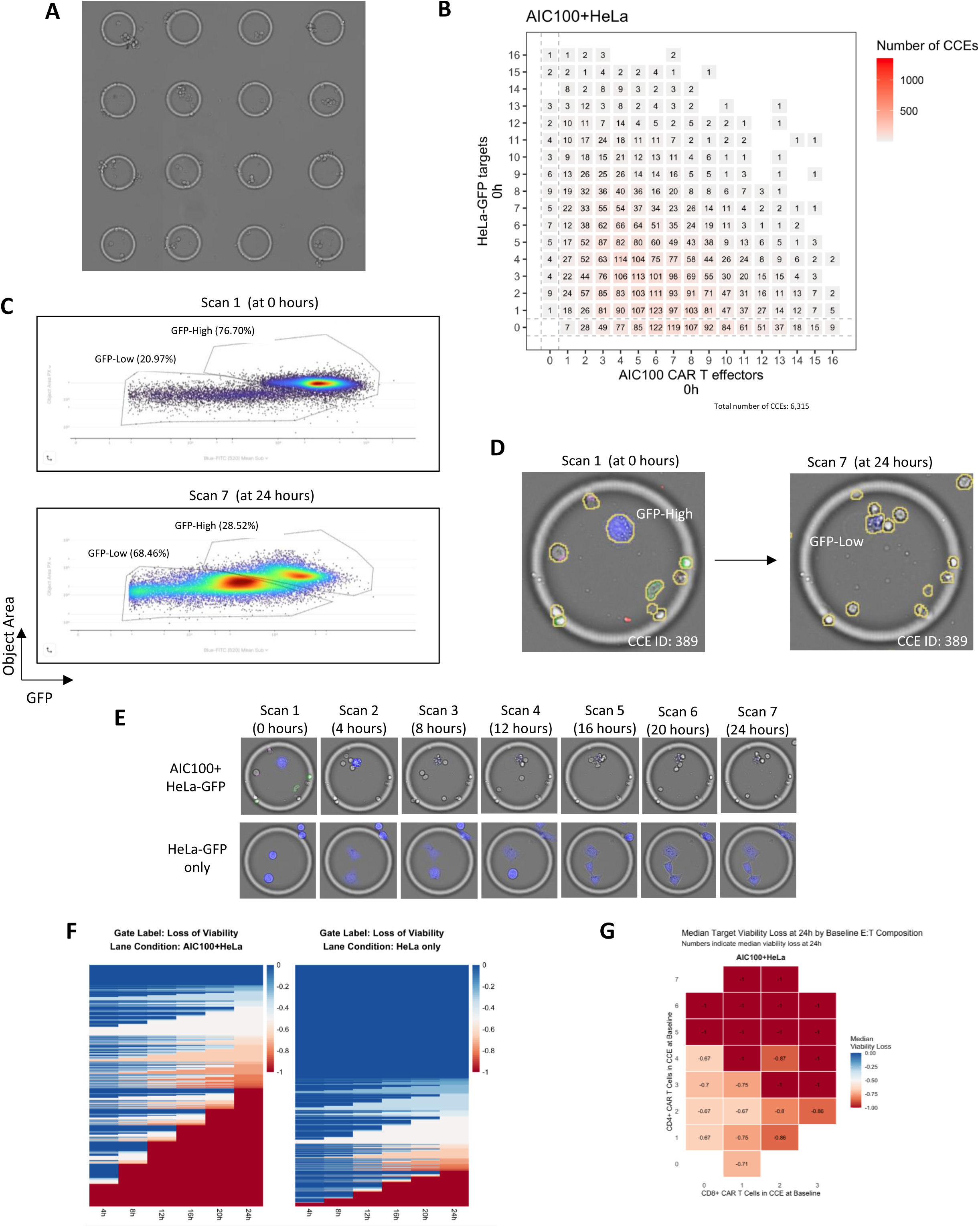
Stain-free cytotoxicity assay using AIC100 CAR-T cells at various E:T ratios and correlating that activity with different T-cell type combinations. (A) Fixed pattern CCE formation arranged in a grid incorporates a stochastic distribution of effectors and targets. Cell identities were not assessed during CCE formation as encapsulation was passive for both effectors and targets. (B) Heatmap depicting the distribution of CCE compositions at baseline (0 hours) for CCEs formed with AIC100 effectors and HeLa-GFP targets, with the number of live AIC100 CAR-T effectors (x-axis) and live HeLa-GFP target cells (y-axis) per CCE shown across a total of 6,315 CCEs. Each tile represents a unique effector-target combination, with color intensity and annotated numerical values reflecting the number of CCEs containing that specific composition. (C) Dot plots depicting the distribution of HeLa-GFP target cells within CCEs based on GFP fluorescence intensity (x-axis) and cell size (y-axis) at two timepoints: 0 hours (top) and 24 hours (bottom). Two distinct target cell subpopulations are resolved by gating: a GFP-high/large population corresponding to live HeLa-GFP target cells, and a GFP-low/small population corresponding to dead or dying target cells that have undergone a reduction in both GFP fluorescence intensity and cell size consistent with the morphological changes associated with cell death. (D) Representative images of a single CCE (ID: 389) at two timepoints, 0 hours and 24 hours, illustrating the transition of HeLa-GFP target cells from the GFP-high/large to the GFP-low/small gate over the course of the experiment. Cell segmentation masks are overlaid on each image, with yellow outlines denoting individually segmented cells. (E) Representative time-lapse brightfield and fluorescence overlay images depicting the contrasting behavior of HeLa-GFP target cells across two conditions over successive imaging timepoints spanning 0 to 24 hours. GFP fluorescence (blue) identifies HeLa-GFP target cells throughout the experiment. Anti-CD4-RY586 and anti-CD8-Alexa Fluor 405 fluorescence, imaged only during the initial timepoint, identifies CD4^+^ and CD8^+^ AIC100 CAR-T effectors, respectively. The loss of GFP fluorescence and decrease in size of HeLa-GFP targets indicates cell death. (F) Longitudinal heatmaps depicting the proportion of live HeLa-GFP target cells lost since baseline within individual CCEs across two experimental conditions: AIC100 CAR-T effectors co-enclosed with HeLa-GFP targets (left), and HeLa-GFP targets enclosed in the absence of effectors (right). Each row represents a single CCE tracked across 6 post-baseline time points (4 through 24 hours), with color intensity reflecting the magnitude of target cell viability loss from the initial timepoint at 0 hours, ranging from no loss (blue) to complete target cell death (red). (G) Heatmap depicting median HeLa-GFP target cell viability loss at 24 hours as a function of baseline effector composition within individual CCEs for the AIC100+HeLa condition. The x-axis and y-axis represent the number of live CD8^+^ and CD4^+^ cells per CCE at baseline, respectively, with each tile corresponding to a distinct effector cell combination. Color intensityreflects the magnitude of median target cell viability loss, with deeper red shading indicating greater cytotoxicity. Numerical values within each tile denote the median fractional viability loss from 0 hours to 24 hours.

As AIC100 effectors were generated from a heterogeneous CD4^+^/CD8^+^ population, anti- CD4 and anti-CD8 fluorescence was imaged at the initial timepoint to enumerate effector subset composition per CCE. Fixed pattern formation yielded a broad range of CD4^+^:CD8^+^ ratios (Figure S6E), with target cell killing scaling positively with CD8^+^ effector number and CCEs containing both CD8^+^ and CD4^+^ cells exhibiting greater killing than those with CD8^+^ effectors alone (Figure 6G). These observations corroborate prior studies demonstrating a helper role for CD4^+^ T lymphocytes in CTL-mediated cytotoxicity^36,40^ and extend these findings to a solid tumor CAR-T context at single-CCE resolution.

## Discussion

Understanding the mechanisms that govern CAR-T cell-mediated cytotoxicity at the single-cell level remains a critical challenge in the development of next-generation cellular immunotherapies. Conventional bulk co-culture assays inherently obscure the functional heterogeneity of individual effector-target interactions, masking key determinants of cytotoxic efficacy such as effector motility, serial killing capacity, and cooperative T cell behavior. Here, we demonstrate that the Cellanome platform overcomes these limitations by enabling the isolation, longitudinal tracking, and high-throughput analysis of defined effector-target combinations within individual CCEs, revealing cellular behaviors that are fundamentally inaccessible to conventional methodologies.

A central finding is the profound functional heterogeneity within CAR-T cell populations. While serial killing by CTLs has been previously reported, prior studies relied on manual tracking of a few dozen cells or indirect mathematical modeling^33,41^, and bulk conditions make it difficult to disentangle cell-intrinsic from cell-extrinsic determinants^42^. Here, 56% of individual effectors engaged in serial killing, a finding derived from direct observations of thousands of effector-target combinations, enabling robust statistical analyses of killing heterogeneity. Analyses at this scale also revealed effector motility and effector-target contact dynamics as independent correlates of cytotoxic potency. Spatiotemporal analysis further showed that direct cell-cell contact strongly predicts target cell death: Annexin V acquisition among proximal targets peaked at approximately 6 hours, after which effectors disengaged and migrated on, while targets that remained distant from effectors throughout stayed Annexin V-negative. Together, these dynamics provide mechanistic evidence linking physical effector-target interactions to cytotoxic outcomes at a resolution unattainable in bulk assays.

When multiple CAR-T cells were co-enclosed with targets, cytotoxic efficacy scaled positively with effector number across both the anti-CD19 and AIC100 CAR-T models, with a substantial proportion of CCEs reaching maximal target cell death within the first 6–12 hours, consistent with recent literature demonstrating that optimal tumor cell killing requires the cumulative cytolytic activities of multiple CTLs^33,34^. The temporal heterogeneity in killing across the CCE population, with individual CCEs reaching maximal cytotoxicity at distinct timepoints, correlated with effector-target engagement dynamics and underscores the importance of resolving cytotoxic activity at the level of individual cellular interactions rather than averaging across bulk populations.

The Cellanome platform further enabled systematic interrogation of cooperative interactions between CD4^+^ T helper cells and CD8^+^ CTLs during the effector phase of tumor cell killing, corroborating *in vivo* studies implicating CD4^+^ T cells as active participants in CTL-mediated tumor clearance^36,40^. Importantly, the CD4^+^ T cells used here are untransduced and do not express the anti-CD19 CAR, ruling out a direct cytotoxic contribution and implicating a contact- or paracrine-dependent cooperative mechanism operating locally between co-enclosed cells. Furthermore, the magnitude of CD4^+^-mediated enhancement of CD8^+^ cytotoxicity scaled dose-dependently with CD4^+^ cell number per CCE, a quantitative relationship not previously resolvable at single-cell resolution, providing a new framework for understanding how the local stoichiometry of CD4^+^ and CD8^+^ T cells influences cytotoxic outcomes, with direct implications for the rational design of CAR-T cell products with optimized CD4^+^:CD8^+^ ratios.

Extending the platform to a solid tumor model using AIC100 anti-ICAM-1 CAR-T cells and HeLa-GFP targets further demonstrated the versatility and generalizability of the CCE-based cytotoxicity assay. Individual AIC100 effectors failed to kill HeLa-GFP targets effectively, whereas multi-effector CCEs exhibited robust and progressive killing consistent with the requirement for coordinated multi-CTL activity in solid tumor elimination^33,35^. Critically, by physically isolating thousands of CCEs containing precisely defined and variable effector numbers, our platform directly measures how killing efficacy scales with effector abundance at single-CCE resolution, a level of quantitative resolution not achievable in prior 3D organotypic or *in vivo* models. The stain-free nature of target cell death quantification in this model, based on loss of GFP fluorescence and changes in cell morphology, further highlights the adaptability of the platform to diverse experimental contexts.

Collectively, these findings establish our platform as a powerful tool for high-resolution, high-throughput characterization of CAR-T cell-mediated cytotoxicity. The ability to longitudinally track thousands of individually isolated effector-target combinations, and to directly correlate cytotoxic potency with defined effector compositions at the single-cell level, has direct translational relevance to the rational optimization of CAR-T cell therapies. Future studies investigating additional determinants of CAR-T cell function, including tumor microenvironment factors, antigen density, and CAR construct design, will further expand our mechanistic understanding of adoptive cellular immunotherapy.

## Methods

### Cell culture

TALL-104 cells (ATCC) were cultured in RPMI-1640 medium supplemented with 20% FBS, 1% PenStrep, and 200 Units/ml recombinant human IL-2. K562 cells (ATCC) were cultured in IMDM medium supplemented with 10% FBS and 1% PenStrep. HeLa-GFP and NALM6- Luc/GFP (Creative Biogene) were cultured in RPMI-1640 medium supplemented with 10% FBS and 1% PenStrep, with NALM6-Luc/GFP medium additionally supplemented with 0.5 μg/mL puromycin. All cells were maintained in a humidified incubator at 37 °C with 5% CO₂..

### T-cell isolation, activation, and culture

AIC100 CAR-T cells were activated with ImmunoCult™ Human CD3/CD28 T Cell Activator (STEMCELL Technologies) and cultured in TexMACS medium supplemented with 10% Human AB Serum, 1% PenStrep, 12.5 ng/mL IL-7, and 12.5 ng/mL IL-15. CD4⁺ and CD8⁺ cells were isolated from bulk anti-CD19 CAR-T cells (BPS Bioscience) and donor matched untransduced T cells (BPS Bioscience) using the MojoSort™ Human CD4 T Cell Isolation Kit and the MojoSort™ Human CD8 T-Cell Isolation Kit (BioLegend), respectively. After isolation, cells were activated with ImmunoCult™ Human CD3/CD28 T-Cell Activator (STEMCELL Technologies) and cultured in T cell medium (BPS Bioscience) with recombinant human IL-2 (200 Units/ml, Miltenyi Biotec). Cells were maintained in a humidified incubator at 37 °C with 5% CO₂.

### Cell labeling

Cells were labeled with indicated fluorescent antibodies at a 1:20 dilution in their respective cell culture medium and incubated in the dark at room temperature for 30 minutes. Cells were then washed and resuspended in fresh medium. For CellTrace CFSE labeling, K562 cells were washed with 1x PBS, resuspended to a concentration of 1x10^6^ cells/mL in a 1 µM CellTrace CFSE staining solution (diluted in 1x PBS), and incubated at 37°C for 20 minutes in the dark. After staining, cells were washed with culture medium, resuspended to a concentration of 1x10^6^ cells/mL in culture medium, and incubated at 37°C for 10 minutes in the dark.

### Cell loading for CCE formation

For CCE formation targeting single unfluorescent effectors, 1.68×10⁵ GFP-negative effectors were combined with 1×10⁶ GFP-positive targets in 70 µL of culture medium. Cells were then loaded into the Cellanome instrument, mixed with hydrogel precursor, and introduced into the flow cell. CCEs were formed around GFP-negative effectors, with GFP-positive targets passively co-enclosed. In negative control lanes, 1.68×10⁵ GFP-negative targets were used as substitutes for effectors, and CCEs were formed around these GFP-negative targets.

For CCE formation targeting single fluorescent effectors, effectors were first labeled with anti-CD4-RY586 or anti-CD8-RY586 antibodies (as indicated). Briefly, 1×10⁶ cells were resuspended in 100 µL of culture medium supplemented with antibodies at a 1:20 dilution, incubated at room temperature for 15 minutes, and washed once with 1 mL of culture medium. Labeled effectors (1.68×10⁵ cells) were then combined with 1×10⁶ unlabeled targets in 70 uL and loaded into the Cellanome instrument for CCE formation as described. CCEs were formed around individual RY586-positive cells, with unlabeled cells passively co-enclosed. For the negative control, NALM6-GFP targets were labeled with anti-CD19-RY586 antibodies using the same protocol, and 1.68×10⁵ labeled targets were combined with 1×10⁶ unlabeled targets for CCE formation. For fixed-pattern CCE formation, 1×10⁶ effectors were combined with 1×10⁶ targets in 70 µL for CCE formation. The negative control consisted of 1×10⁶ targets alone, prepared in 70 µL.

### Cytotoxicity Assays

For cytotoxicity assays, harvested effector cells were washed and resuspended in the respective target cell medium. After CCE formation, cell culture medium with or without APC-Annexin V supplemented at a 1:200 dilution was applied to the lane to wash away any unpolymerized hydrogel precursor. Time-lapse imaging was performed at the indicated intervals and with the indicated fluorescence channels.

### Deep-learning based segmentation and feature extraction of cells

Detection of cell positions and image-based feature extraction was achieved using two in-house deep learning models, trained on an in-house dataset consisting of ∼3 million manually annotated object masks from ∼15,000 images. Cells were identified for CCE formation, either in brightfield or fluorescent channels, using bounding boxes from the fast detector model YOLOv5^43^, implemented with Ultralytics (www.ultralytics.com). For in-depth image analysis used to classify cell identities and states, we used masks from the instance segmentation model Mask R-CNN^44^, implemented with PyTorch (pytorch.org). Cells were segmented in brightfield, and other metrics such as shape and intensity computed from each of those masks. Fluorescent signal was computed per channel from the mean intensity of all pixels that comprise an object instance, with background subtraction performed by further subtracting the mean intensity of all non-foreground pixels within that CCE. All cells were given unique identifiers, and were associated with CCEs which were also given unique identifiers, that are registered over time using fiducials. All processing steps were implemented in an image analysis pipeline written in Python and Nextflow^45^.

### User-in-the-loop UI for identification of cell populations

A cloud-based user interface (UI) enables users to visualize the distribution across all cells of the features computed by the image analysis pipeline. Thereby, in a 1-dimensional or 2-dimensional plot, users can select subpopulations of cells or “gates” based on criteria of cell morphology and fluorescence intensity, and create hierarchically defined populations for further analysis, using functionality that displays the image of the cell behind each data point to refine and correct the boundaries of these populations. This interface allowed gates to be tailored to the distributions of each experiment, while supporting the creation of standardized gates used for further analysis across all datasets shown: Live Target cells, Dead Target cells, Live Effector cells, and Dead Effector cells. Per-CCE counts for each gate are used for filtering steps, to calculate E:T (Effector:Target) ratios for each CCE, and finally to compute cell viability outcomes for each CCE that are based on cell identity memberships.

### Longitudinal analysis of target viability loss in CCEs

Per-CCE counts of gated objects at each timepoint are used to compute longitudinal trends of target viability loss. This is used instead of relying on counts of Dead Target-gated cells over time, which are subject to false positive errors when dead cells fragment and are segmented as multiple cells, or false negative errors when dead cells have completely lysed or eventually lose Annexin V fluorescence and are no longer segmented or gated as dead target cells. By anchoring viability on known counts of Live Target cells at baseline, the timepoint at which cells are most intact and viable, we used changes in the count of Live Target cells at each subsequent timepoint in that CCE to monitor the loss of viable target cells. Cells could lose viability either by changing membership from the Live Target gate to Dead Target gate by update of Annexin V signal, or by no longer being segmented at all, if the cell completely lysed or its morphology became unrecognizable as a cell. In the case of GFP-expressing HeLa cells, a similar strategy was employed by anchoring the analysis on viability loss tracked as a transition from a GFP-high gate to a GFP-low gate as target cells lost viability. Target viability loss over time, computed in this manner assuming a monotonically decreasing trend, enabled a more accurate quantification of cytotoxicity that was more robust to segmentation and gating errors amid the complexities of closely interacting cells with deteriorating morphology.

### Quantification of cell-cell interactions and effector motility over time

Target-effector proximity, and its correlation to other metrics such as Annexin V uptake, was computed using the known positions and identities of each cell in a CCE at any given timepoint. This analysis was filtered to CCEs with only one cell gated as an effector, and 2 or more target cells, at all timepoints. The plot for each timepoint was computed with a single data point for every target cell in these CCEs, showing the Euclidean distance between that target cell and the effector cell in the same CCE on the x axis, and the background-subtracted mean Annexin V signal on the y-axis of the same target cell at that timepoint. This shows the distribution of proximal versus distant target-effector interactions and their correlation with apoptotic processes, and dividing this data into quadrants allowed us to track overall population shifts over time. The same analysis was performed for the control lane, nominating a single random cell as a pseudo-effector in each CCE in the absence of actual effector cells, to compute the Euclidean distance between each target cell and that nominated cell. This enabled a comparison of the distribution of target cells between the proximal/distant and low/high-Annexin V quadrants between the co-cultured and control conditions at each timepoint. Effector motility was also computed only for CCEs with only one cell gated as an effector, and 2 or more target cells, at all timepoints, in order to track the movement of effector cells that had access to multiple target cells. Positional coordinates for each cell are referenced against fiducials to generate a global set of coordinates that are consistent over time and corrected for stage movement errors. These global positional coordinates can be used to calculate the Euclidean distance travelled by the effector cell at each timepoint since its position in the previous timepoint, and summed across all timepoints to arrive at the total path length travelled by that effector. This leads to a distribution of path lengths across all effector cells, which was divided into tertiles to define high-motility (>72 µm), medium-motility (37-72 µm), or low-motility (<37 µm) groups of effectors. This functional property of each effector could then be correlated with other metrics such as cytotoxicity.

## Supplementary Figures

**Supplementary Figure 1.**
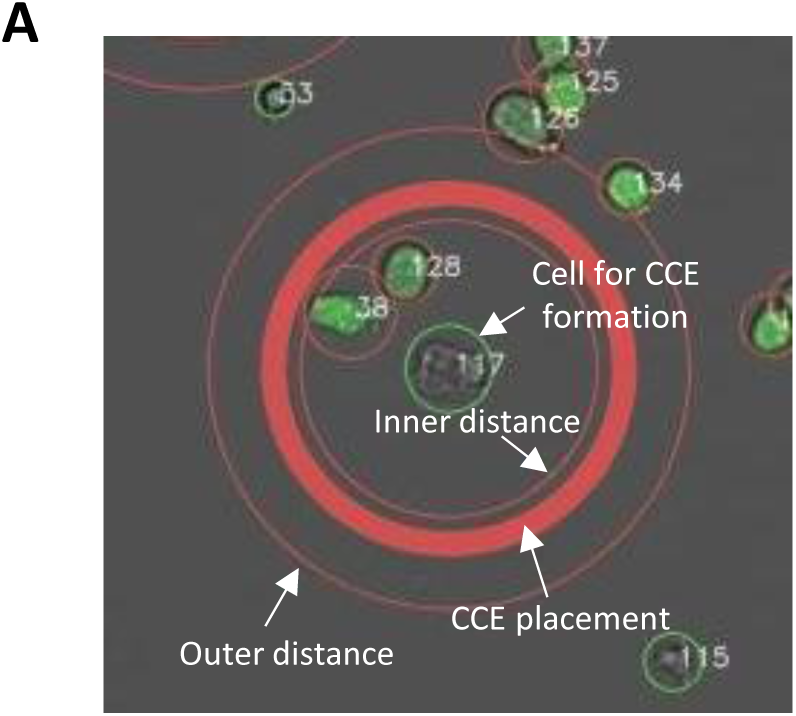
(A) Representative fluorescence image illustrating the spatial parameters governing targeted CCE formation on the Cellanome platform. A cell selected for CCE formation is shown at the center, flanked by two concentric rings that define the user-specified spatial constraints for CCE wall placement. The inner ring delineates the minimum allowable distance between the target cell and the inner CCE wall, ensuring that the target cell is fully enclosed within the CCE without direct contact with the hydrogel matrix. The outer ring defines the maximum allowable distance for CCE wall placement, beyond which CCE formation will not occur. Together, these two parameters define an annular zone within which the CCE wall must be positioned for successful encapsulation.

**Supplementary Figure 2.**
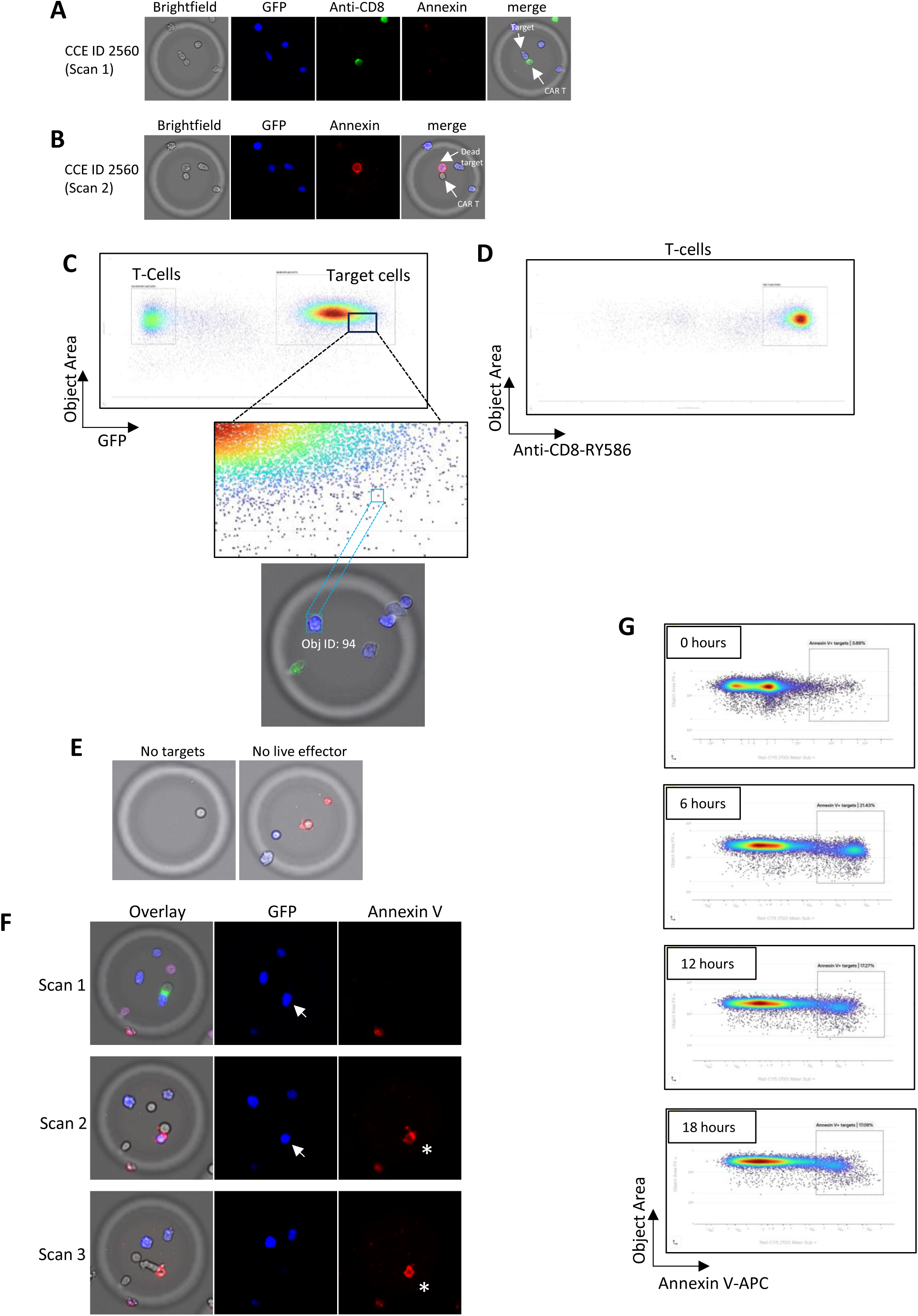
(A) Images of a single CCE (ID: 2560) at the initial imaging timepoint, showing a co-enclosed CD8^+^ CAR-T effector and NALM6-GFP target cells. From left to right: brightfield image, GFP fluorescence (blue), anti-CD8-RY586 fluorescence (green), APC-Annexin V fluorescence (red), and a merged brightfield-fluorescence overlay. The merged image confirms the presence of a single GFP-negative CD8^+^ CAR-T effector alongside multiple GFP-positive NALM6-GFP target cells within the CCE, with minimal APC-Annexin V signal, indicating that target cells are viable at baseline. (B) Images of the same CCE shown in (A) at the 6 hour imaging timepoint. From left to right: brightfield image, individual fluorescence channels displaying GFP (blue) and APC-Annexin V (red) signals, and a merged brightfield-fluorescence overlay. In contrast to the initial timepoint, the target engaged with the effector acquired strong APC-Annexin V fluorescence, indicative of effector-mediated induction of apoptosis. (C) Representative illustration of the interactive dot plot analysis feature within the Cellanome cloud-based analysis platform, demonstrating the ability to link quantitative single-cell measurements to their corresponding microscopy images. The top panel shows a dot plot in which two distinct cell populations are resolved based on measured parameters: T-cells (left gate) and NALM6-GFP target cells (right gate), separated by differences in fluorescence intensity and cell size. The middle panel displays a zoomed-in view of the target cell gate. A single data point within this gate is selected. The bottom panel displays the corresponding image of the individual cell linked to the selected data point, identified as Object ID 94, within its encapsulating CCE. (D) Dot plot depicting anti-CD8-RY586 fluorescence (x-axis) and cell size (y-axis, object area in pixels) of cells within the “T-cells” gate shown in Figure S2C, confirming the cells within this gate are CD8^+^ CAR-T effectors. (E) Representative images of two CCEs excluded from downstream cytotoxicity analyses due to uninformative cell compositions. The first CCE (left) contains a single GFP-negative CAR-T effector in the absence of any NALM6-GFP target cells, precluding assessment of cytotoxic activity. The second CCE (right) contains NALM6-GFP target cells (blue) alongside a non-viable Annexin V-positive effector (red). Exclusion of such CCEs from downstream analysis reduces background signal and ensures that cytotoxic activity is quantified only in CCEs containing both viable effectors and target cells. (F) Representative images of a single CCE across three successive imaging timepoints (top to bottom), illustrating the progressive loss of GFP fluorescence following CAR-T cell-mediated target cell death. Merged brightfield-fluorescence overlay (left) with individual GFP (blue) and APC-Annexin V (red) fluorescence channel images (right) are shown for each timepoint. White arrow indicates a GFP-positive NALM6-GFP target cell engaged with the CAR-T effector, and asterisk indicates Annexin V staining for this target cell. In the first timepoint, the indicated target is GFP-positive and Annexin V-negative, and transitions to the second timepoint where the GFP and Annexin V are double positive. By the last timepoint, this target cell has lost all GFP fluorescence and is only Annexin V-positive. (G) Dot plots depicting the proportion of NALM6-GFP target cells that are also Annexin V positive across four successive imaging timepoints (Timepoints 0-18 hours). For each timepoint, APC-Annexin V fluorescence (x-axis) is plotted against cell size (y-axis, object area in pixels). The apparent decline in Annexin V-positive targets does not reflect a true reduction in target cell death, but rather an artifact arising from the progressive loss of GFP fluorescence in late-stage dead target cells. As dead target cells transition from GFP-positive/Annexin V-positive to GFP-negative/Annexin V-positive over time, they exit the GFP-positive target cell gate entirely and are no longer counted as Annexin V-positive target cells in subsequent timepoints.

**Supplementary Figure 3.**
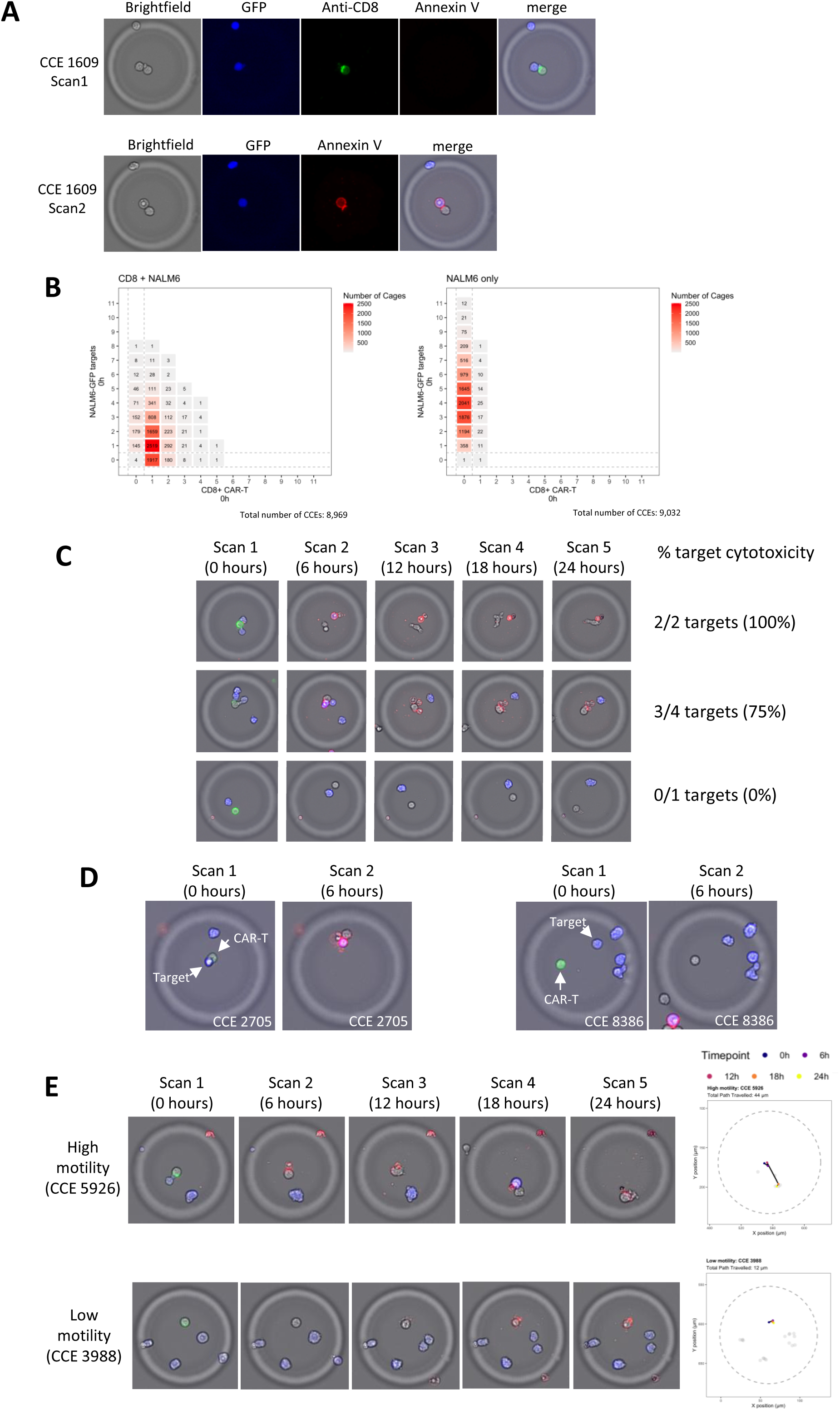
(A) Images of a CCE ID: 1609 at the initial imaging timepoint (top) and at 6 hours (bottom) showing individual fluorescence signals. Top panels from left to right; brightfield image, GFP fluorescence (blue), Anti-CD8 (green), APC-Annexin V fluorescence (red), and a merged brightfield-fluorescence overlay. Images confirm the identity of the GFP-negative CD8^+^ CAR-T effector and GFP-positive target, with minimal APC-Annexin V signal, indicating viability of both cells at baseline. Bottom panels from left to right; brightfield image, GFP (blue) and APC-Annexin V (red) signals, and a merged brightfield-fluorescence overlay. In contrast to the initial timepoint, the target has acquired strong APC-Annexin V fluorescence at Scan 2, indicative of effector-mediated induction of apoptosis. (B) Heatmaps depicting the distribution of CCE compositions at baseline (0 hours) for two experimental conditions: CD8^+^ CAR-T effectors co-enclosed with NALM6-GFP targets (left, total CCEs = 8,969) and NALM6-GFP targets enclosed in the absence of effectors (right, total CCEs = 9,032). For each condition, the number of CD8^+^ CAR-T cells (x-axis) and NALM6-GFP target cells (y-axis) per CCE are shown, with each tile representing a unique cell combination. Color intensity and annotated numerical values reflect the number of CCEs containing that specific composition. Dashed lines delineate CCEs containing zero effectors or zero target cells. (C) Example time-lapse images depicting CAR-T cell-mediated cytotoxic activity within individual CCEs that enclosed various E:T ratios resulting in different proportion of target cell death. GFP fluorescence (blue) marks live NALM6-GFP target cells, green fluorescence identifies the CAR-T effector at the initial timepoint only, and APC-Annexin V fluorescence (red) indicates target cell death. CCE-level analysis enables the quantification of percentage of target cell death for each CCE. (D) Representative time-lapse fluorescence overlay images from two CCEs illustrating the relationship between effector-target proximity and cytotoxic outcome. GFP fluorescence (blue) marks live NALM6-GFP target cells, green fluorescence identifies the GFP-negative CAR-T effector, and APC-Annexin V fluorescence (red) indicates cell death. In CCE 2705 (left panels), a CAR-T effector is observed in direct contact with a target cell at the initial timepoint. In the subsequent imaging timepoint, the engaged target cell acquires strong Annexin V fluorescence, indicative of effector-mediated cytotoxicity, while neighboring target cells that were not in direct contact with the effector remain Annexin V-negative. In contrast, in CCE 8386 (right panels), the CAR-T effector is spatially distant from the target cell population at the initial timepoint. Consistent with the absence of direct effector-target contact, target cells in this CCE remain Annexin V-negative in the subsequent imaging timepoint, indicating sustained viability. (E) Example images of high-motility and low-motility effectors. Images of a CCE with a highly motile effector (CCE ID: 5926) from all timepoints, observed initially engaging a target cell in close proximity and then killing it (as evidenced by uptake of Annexin V and loss of GFP signal), and then moving to a second target cell at 18 hours and eventually killing it as well. The trace of this effector’s path to the CCE is indicated on the right, with colored markers indicating its position at each timepoint, and grey markers indicating the positions of the target cells at all timepoints. Images of a CCE with a low-motility effector (CCE ID: 3988) from all timepoints, observed remaining in the same position throughout all timepoints (as visualized with its path trace on the right), despite the presence of multiple live target cells in the same CCE. This low-motility effector eventually takes up Annexin V and dies without any impact on the viability of its co-enclosed target cells, in contrast to the highly motile effector above.

**Supplementary Figure 4.**
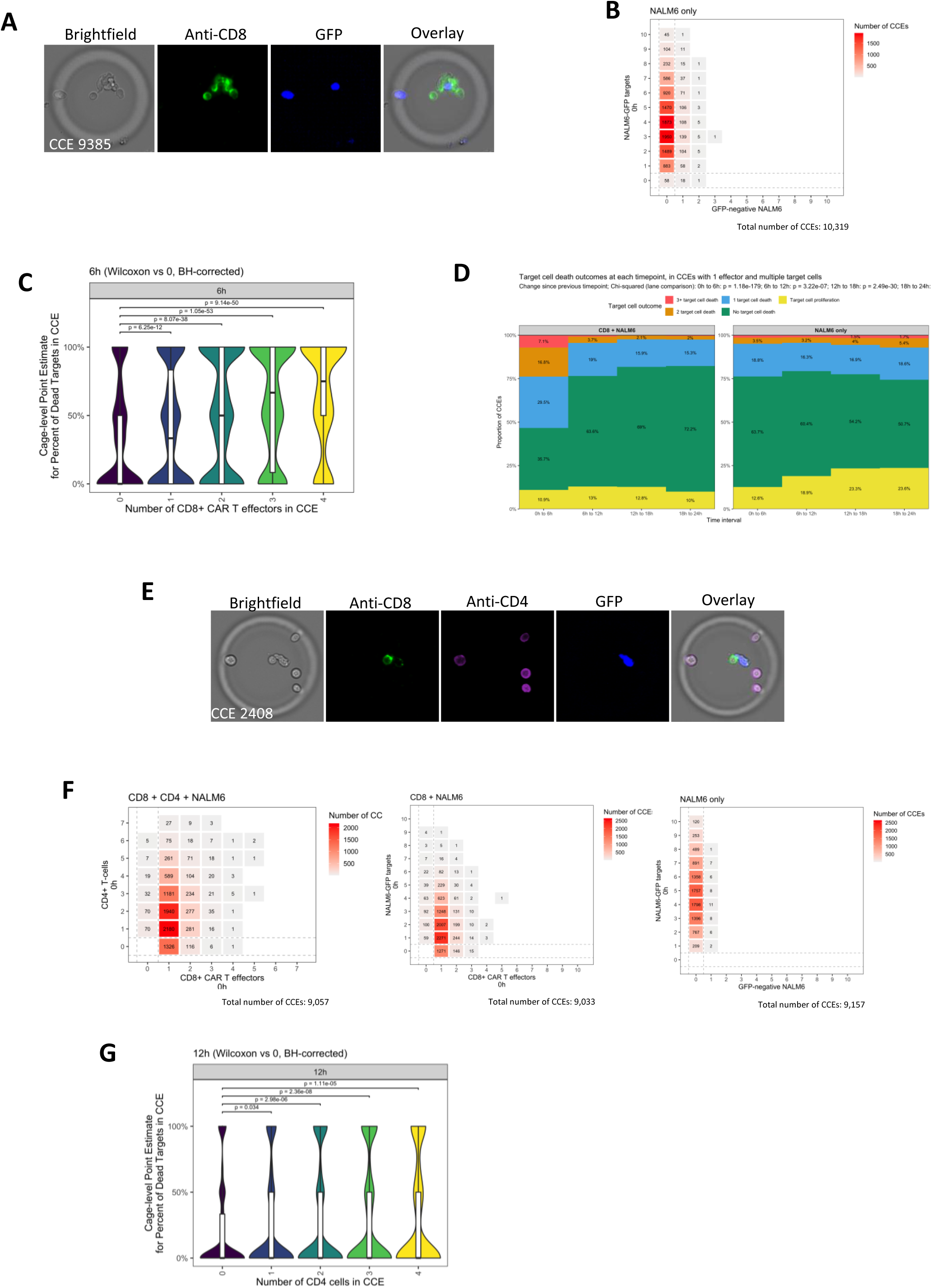
(A) Images of a single CCE (ID: 9385) at the initial imaging timepoint. From left to right: brightfield image, anti-CD8-RY586 fluorescence (green), GFP fluorescence (blue), and a merged brightfield-fluorescence overlay. Multiple GFP-negative CD8^+^ CAR-T effectors, identified by anti-CD8-RY586 fluorescence, actively engaging GFP-positive NALM6-GFP target cells within the CCE. (B) Heatmaps depicting the distribution of CCE compositions at baseline (0 hours) for the control condition: NALM6-GFP targets enclosed in the absence of effectors (total CCEs = 10,319). The number of GFP-negative NALM6 cells (x-axis) and NALM6-GFP target cells (y-axis) per CCE are shown, with each tile representing a unique cell combination. Color intensity and annotated numerical values reflect the number of CCEs containing that specific composition. Dashed lines delineate CCEs containing zero effectors or zero target cells. (C) Violin plots showing the distribution of target cell death (the percentage of target cells within a CCE that are Annexin V-positive) at 6 hours, in different subsets of CCEs within the co-enclosed lane. Each violin represents the group of CCEs with the specified number of effector cells in the CCE, with Benjamini-Hochberg corrected p-values from the Wilcoxon test displayed for each subset compared against the 0-effector CCEs within this lane. These depict an increasing trend of target cell death with increasing effector count. (D) Stacked bar charts depicting the distribution of target cell death outcomes within individual CCEs encapsulating a single CD8^+^ CAR-T effector and multiple NALM6-GFP target cells (left) compared to NALM6-GFP targets enclosed in the absence of effectors (right). The proportion of CCEs exhibiting each outcome category, ranging from target cell proliferation and no target cell death to the loss of 1, 2, or 3 or more target cells, was quantified using the change in live target cell count from 0 hours to each timepoint. A chi-squared test was computed on the distribution of categories between the two lanes. (E) Images of a single CCE (ID: 2408) at the initial imaging timepoint. From left to right: brightfield image, anti-CD8-RY586 fluorescence (green), anti-CD4-Alexa Fluor 405 fluorescence (violet), GFP fluorescence (blue), and a merged brightfield-fluorescence overlay. Fluorescence images confirm the co-encapsulation of a single CD8^+^ CAR-T effector, multiple CD4^+^ T helper cells, and GFP-positive NALM6-GFP target cells within the same CCE. (F) Heatmaps depicting the distribution of CCE compositions at baseline (0 hours) for three experimental conditions (Left) CD8^+^ CAR-T effectors co-enclosed with both CD4^+^ untransduced donor-matched T-cells and NALM6-GFP targets (total CCEs = 9,057) showing the number of CD8^+^ CAR-T cells (x-axis) and CD4^+^ T-cells (y-axis) per CCE. (Middle heatmap) CD8^+^ CAR-T effectors co-enclosed with NALM6-GFP targets without CD4^+^ untransduced cells (total CCEs = 9,033), the number of CD8^+^ CAR-T cells (x-axis) and NALM6-GFP target cells (y-axis) per CCE are shown. (Right heatmap) NALM6-GFP targets enclosed in the absence of effectors (total CCEs = 9,157),, the number of anti-CD19-RY586 stained NALM6-GFP cells (x-axis) and unstained NALM6-GFP target cells (y-axis) per CCE are shown. For all heatmaps, each tile represents a unique cell combination. Color intensity and annotated numerical values reflect the number of CCEs containing that specific composition. Dashed lines delineate CCEs containing zero effectors or zero target cells. (G) Violin plots showing the distribution of target cell death (the percentage of target cells within a CCE that are Annexin V-positive) at 6 hours, in different subsets of CCEs within the co-enclosed lane. Each violin represents the group of CCEs with the specified number of CD4^+^ untransduced donor-matched T-cells in the CCE, with Benjamini-Hochberg corrected p-values from the Wilcoxon test displayed for each subset compared against the 0-effector CCEs within this lane.

**Supplementary Figure 5.**
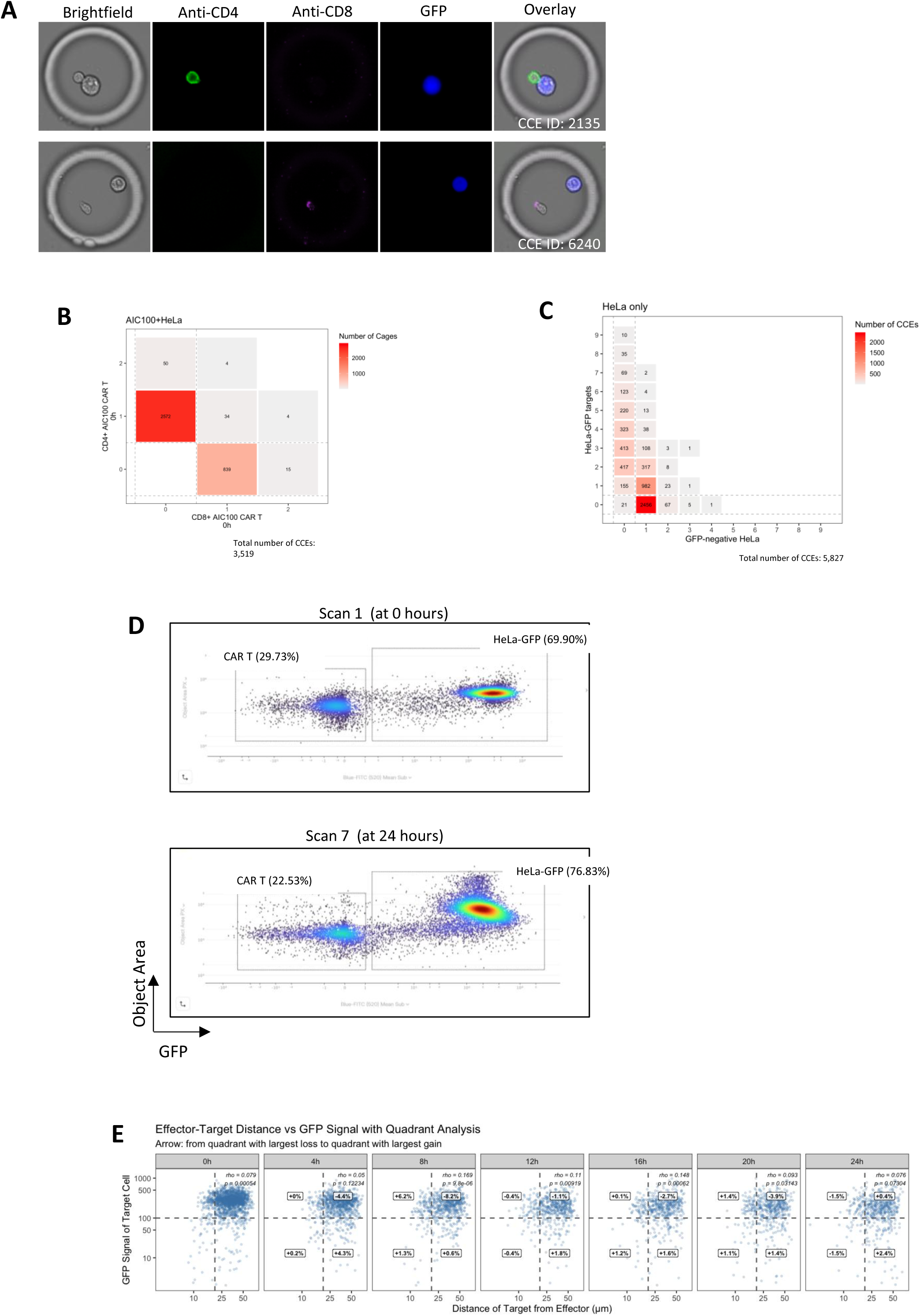
(A) Images of two individual CCEs at the initial imaging timepoint, illustrating the selective encapsulation of distinct AIC100 CAR-T effector subtypes. From left to right: brightfield image, anti-CD4-RY586 fluorescence (green), anti-CD8-Alexa Fluor 405 fluorescence (violet), GFP fluorescence (blue) signals, and a merged brightfield-fluorescence overlay. In CCE 2135 (top row), a single CD4^+^ CAR-T effector, identified by anti-CD4-RY586 fluorescence, is co-enclosed with a GFP-positive HeLa-GFP target cell, with no detectable CD8^+^ signal, confirming the presence of a single CD4^+^ effector in isolation. In CCE 6240 (bottom row), a single CD8^+^ CAR-T effector, identified by anti-CD8-Alexa Fluor 405 fluorescence, is co-enclosed with a GFP-positive HeLa-GFP target cell, with no detectable CD4^+^ signal, confirming the presence of a single CD8^+^ effector in isolation. (B) Heatmap depicting the distribution of CCE compositions at baseline (0 hours) for the condition where AIC100 CAR-T effectors were co-enclosed with HeLa-GFP targets (total CCEs = 3,519) showing the number of CD8^+^ AIC100 CAR-T effectors (x-axis) and CD4^+^ AIC100 CAR-T effectors (y-axis) per CCE. Each tile represents a unique cell combination; color intensity and annotated numerical values reflect the number of CCEs containing that specific composition. Dashed lines delineate CCEs containing zero effectors or zero target cells. (C) Heatmap depicting the distribution of CCE compositions at baseline (0 hours) for the control condition: HeLa-GFP targets enclosed in the absence of effectors (total CCEs = 5,827) showing the number of GFP-negative HeLa cells (x-axis) and HeLa-GFP target cells (y-axis) per CCE. Each tile represents a unique cell combination; color intensity and annotated numerical values reflect the number of CCEs containing that specific composition. Dashed lines delineate CCEs containing zero effectors or zero target cells. (D) Dot plots depicting the distribution of enclosed cells within CCEs based on GFP fluorescence intensity (x-axis) and cell size (y-axis) at the 0 hour (top) and 24 hour (bottom) timepoints. Two distinct cell populations are resolved by gating: GFP-negative AIC100 CAR-T effectors (left gate) and GFP-positive HeLa-GFP target cells (right gate), separated based on differences in GFP fluorescence intensity and cell size. Each data point represents an individual cell. (E) Scatter plots depicting the relationship between effector-target distance (x-axis, µm) and GFP signal of individual HeLa-GFP target cells (y-axis) across seven imaging timepoints (0, 4, 8, 12, 16, 20, and 24 hours). Each plot is divided into four quadrants by dashed lines delineating proximal versus distal effector-target distances and GFP-high versus GFP-low target cells, with the latter corresponding to dead target cells that have undergone a reduction in GFP fluorescence.

**Supplementary Figure 6.**
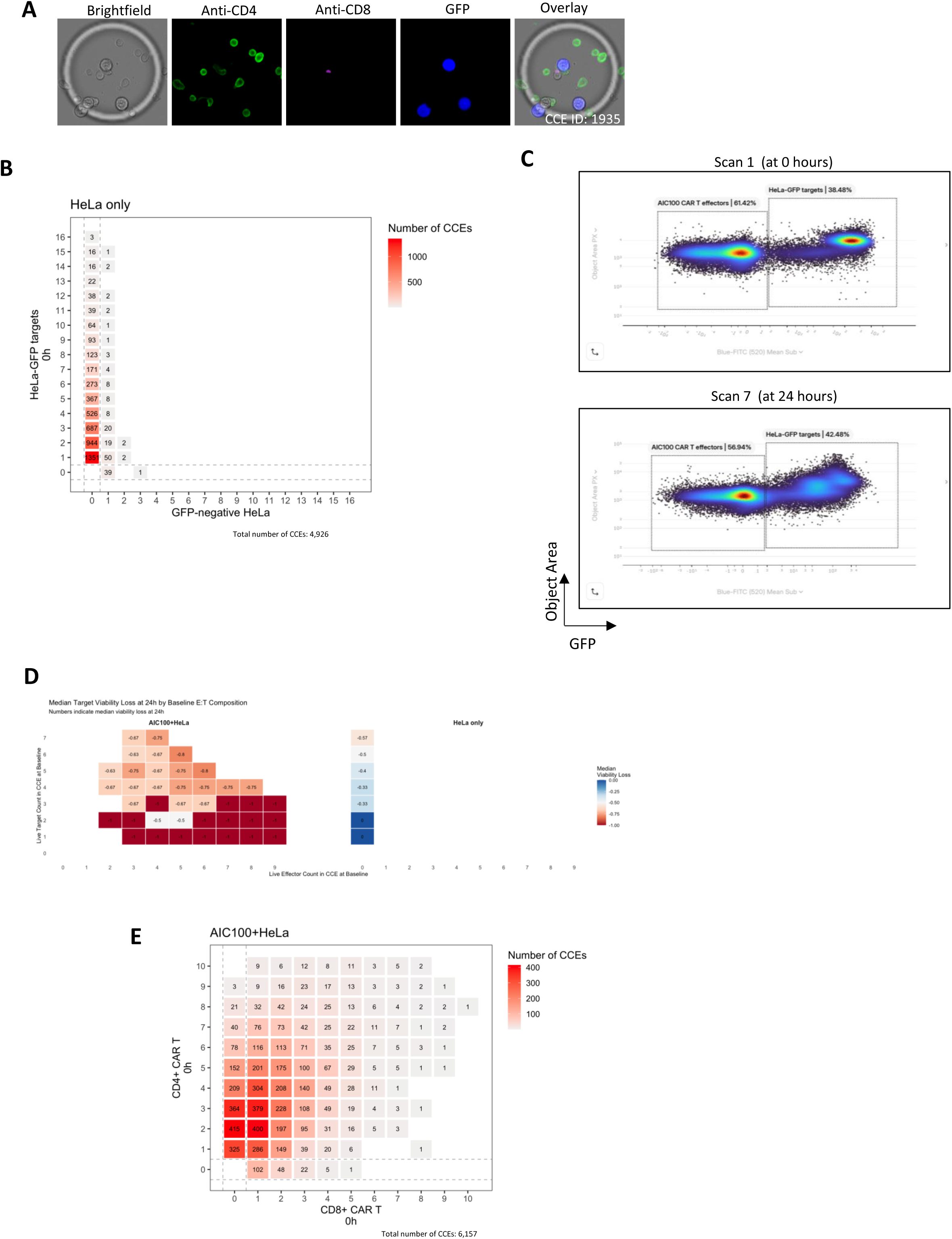
(A) Images of a single CCE at the initial imaging timepoint. From left to right: brightfield image, anti-CD8-RY586 fluorescence (green), anti-CD4-Alexa Fluor 405 fluorescence (violet), GFP fluorescence (blue) signals, and a merged brightfield-fluorescence overlay. Fluorescence images confirm multiple GFP-negative AIC100 CAR-T effectors, predominantly CD8^+^ as identified by anti-CD8-RY586 fluorescence, co-enclosed with a small number of GFP-positive HeLa-GFP target cells and a single CD4^+^ T cell identified by anti-CD4-Alexa Fluor 405 fluorescence. (B) Heatmap depicting the distribution of CCE compositions at baseline (0 hours) for the control condition: HeLa-GFP targets enclosed in the absence of effectors (total CCEs = 4,926) showing the number of GFP-negative HeLa cells (x-axis) and HeLa-GFP target cells (y-axis) per CCE. Each tile represents a unique cell combination; color intensity and annotated numerical values reflect the number of CCEs containing that specific composition. Dashed lines delineate CCEs containing zero effectors or zero target cells. (C) Dot plots depicting the distribution of enclosed cells within CCEs based on GFP fluorescence intensity (x-axis) and cell size (y-axis) at the 0 hour (top) and 24 hour (bottom) time points. Two distinct cell populations are resolved by gating: GFP-negative AIC100 CAR-T effectors (left gate) and GFP-positive HeLa-GFP target cells (right gate), distinguished by differences in GFP fluorescence intensity and cell size. Each data point represents an individual cell. (D) Heatmap depicting median HeLa-GFP target cell viability loss at 24 hours as a function of effector:target ratios within individual CCEs for the AIC100+HeLa condition (left). The x-axis and y-axis represent the number of effector cells and target cells per CCE at baseline (0 hours), respectively, with each tile corresponding to a distinct effector:target combination. Color intensity reflects the magnitude of median target cell viability loss, with deeper red shading indicating greater cytotoxicity. Numerical values within each tile denote the median fractional viability loss from 0 hours to 24 hours. The corresponding median viability loss per target cell count is shown for the control lane, which has 0 effectors for all CCEs (right). (E) Heatmap depicting the distribution of CCE compositions at baseline (0 hours) for the co-enclosed condition: AIC100 CAR-T effectors co-enclosed with HeLa-GFP targets (total CCEs = 6,157) showing the number of CD8^+^ AIC100 CAR-T effectors (x-axis) and CD4^+^ AIC100 CAR-T effectors (y-axis) per CCE. Each tile represents a unique cell combination; color intensity and annotated numerical values reflect the number of CCEs containing that specific composition. Dashed lines delineate CCEs containing zero effectors or zero target cells.

